# Multiple instance learning with spatial transcriptomics for interpretable patient-level predictions: application in glioblastoma

**DOI:** 10.1101/2025.10.13.682206

**Authors:** Simon Grouard, Christian Esposito, Jean El Khoury, Valérie Ducret, Céline Thiriez, Loïc Herpin, Anaïs Chossegros, Caroline Hoffmann, Quentin Bayard, Genevieve Robin, Nicole Tay, Esther Baena, MOSAIC Consortium, Eric Durand, Almudena Espin Perez, Lucas Fidon

## Abstract

Accurate prediction of patient outcomes remains a major challenge in oncology. While recent machine learning (ML) approaches often rely on bulk omics lacking spatial resolution or histology-based multiple instance learning (MIL), spatial transcriptomics (SpT) provides a unique opportunity to capture both molecular content and tissue architecture. However, no generalizable ML framework has yet been established to exploit SpT for patient-level outcomes. We present SpaMIL, a flexible and interpretable MIL framework designed for SpT, with a distillation strategy that enables deployment for hematoxylin and eosin (H&E) slides alone. We evaluate the framework by predicting survival from glioblastoma (GBM) patients, a clinically compelling setting given its aggressiveness with a median survival of only 15 months and the lack of prognostic clinical variables. We analyzed 76 GBM cases from the MOSAIC dataset: 43 with matched SpT, H&E, single-nucleus RNA-seq (scRNA-seq), bulk RNA-seq, and clinical variables, and 33 with H&E for external validation. We developed two main architectures: abMIL, tailored to SpT’s spatial molecular structure, and MabMIL, which distills SpT-derived representations into H&E. Model interpretability was achieved through a Shapley-based framework linking prognostic predictions to cell-type compositions via SpT deconvolution. In benchmarking across the five GBM MOSAIC modalities, SpT-based abMIL achieved unprecedented prognostic accuracy (median C-index: 0.72, standard deviation: 0.04), outperforming all other modalities, including established clinical predictors. PCA and deconvolution-based SpT representations surpassed recent foundation models, suggesting the need for further research on SpT foundation models. Our interpretability analysis highlighted malignant and non-malignant cell subpopulations associated with favorable or poor prognosis, consistent with recent reports. Finally, MabMIL maintained strong performance while enabling H&E-only deployment, with improved condorance index over H&E-only baselines in both internal (0.59 vs. 0.57) and external (0.62 vs. 0.55) cohorts.

## Introduction

Accurate prediction of patient outcomes remains a major challenge in oncology. In glioblastoma, (GBM), the most common primary tumor of the central nervous system, most existing ML models rely on clinical data (Jiang et al. 2021; Chunduru et al. 2022; Chen et al. 2014), bulk RNAseq (Chen, Lu, Wang, et al. 2022) or hematoxylin and eosin (H&E)-stained histopathology slides, which are routinely available in clinical workflows (Jiang et al. 2021; Chunduru et al. 2022; Chen et al. 2014; Redlich et al. 2024) to predict survival (median survival: 15 months; 5-year survival rate: 6.9%). Similar approaches have also been applied in other cancers, including breast, lung, and colorectal cancers (Gross et al. 2024; Filiot et al. 2024).The Multiple Instance Learning (MIL) framework is commonly used to predict patient outcomes from whole-slide images and achieves state-of-the-art performance due to its ability to model survival outcome by aggregating complex information from high-resolution image data^4–13^. Since these images are too large to process directly, they are divided into smaller tiles, each treated as an instance in a bag representing one image. The MIL model learns to assign importance weights to each tile, typically via attention mechanisms, and aggregates them into a single representation used for prediction (Ilse et al. 2018; Saillard et al. 2020). Crucially, the model is trained using only patient-level labels, without requiring annotations for individual tiles.

Recent advances in spatial transcriptomics (SpT) offer new opportunities for modeling cancer heterogeneity. In particular, 10x Genomics Visium enables spatially resolved gene expression profiling across intact tissue sections, preserving local heterogeneity compared to bulk RNAseq and potentially offering more prognostically informative data. We hypothesize that extending the MIL framework, originally developed for H&E, to SpT could substantially improve patient-outcome prediction. GBM provides a particularly challenging and clinically relevant setting to evaluate this approach, given its pronounced intratumoral heterogeneity and the difficulty of accurately modeling patient survival (Hegi et al. 2005a; Gorlia et al. 2008): currently, only a limited set of variables, including patient age, Karnofsky Performance Status (KPS), and O6-methylguanine-DNA methyltransferase (MGMT) promoter methylation status, are consistently associated with survival outcomes under SOC and are routinely used to guide treatment decisions (Chunduru et al. 2022; Chen et al. 2014). Recent work used 22 paired H&E and SpT samples to train a ML model that predicts the cell type composition of each H&E location and used this model on TCGA to associate the presence of clusters of AC-like cells and the colocalization of OPC-like and MES-hypoxia cells, three known GBM malignant cells subtypes (Neftel et al. 2019), with survival. In a more end-to-end approach, graph neural networks (GNNs) applied to SpT data of 130 gliomas including 80 GBM tumors were proposed to improve diagnosis accuracy for gliomas (Ritter et al. 2025).

In this work, we propose SpaMIL, the first end-to-end application of MIL for survival prediction using SpT. We conducted our study using the unprecedentedly large multimodal dataset MOSAIC (MOSAIC Consortium and Hoffmann 2025) comprising 76 GBM samples, including 43 profiled with matched SpT, H&E histology, single-cell RNA sequencing (scRNA-seq), bulk RNA sequencing, and clinical variables, and an additional 33 with H&E for external validation. Leveraging this unique resource, we present the first comprehensive benchmark of survival prediction performance across five data modalities. Our results demonstrated that SpT-based abMIL outperforms all other data modalities for survival prediction. Using our GBM cohort, we systematically investigated optimal spot-level feature representations for survival modeling with SpT and showed that principal component analysis (PCA) and cell type deconvolution features outperformed the more complex representation models scVI (Lopez et al. 2018) and the foundation model scGPT-spatial (Wang et al. 2025). Finally, we assessed model interpretability by applying KernelSHAP to the deconvolution features, ranking cell-type proportions by their contributions to predicted hazard. Malignant and non-malignant cell subpopulations associated with favorable or poor prognosis aligned well with previous work (Neftel et al. 2019; Fidon et al. 2025), supporting that SpT-based MIL captures biologically meaningful tumor–microenvironment interactions rather than dataset artifacts.

Furthermore, we introduce MabMIL, a Multimodal version of abMIL that transfers survival-relevant transcriptomic signals into a histology model. MabMIL distills SpT’s model representation into H&E’s model representation using a teacher-student approach and allows H&E-only inference. Knowledge distillation has previously been applied in computational pathology and more recently in single-cell transcriptomics and multi-omics integration. However, its application to multimodal spatial transcriptomics and histology remains largely unexplored (Filiot et al. 2025; Mann-Krzisnik and Li 2025; Cong et al. 2024; Yu et al. 2025). Our framework enables multimodal models to be trained jointly on SpT and H&E data, while supporting inference using H&E alone, thereby maximizing H&E-only performance while maintaining compatibility with routine clinical data. Although we focused on survival prediction in GBM, our methodology is readily generalizable to other tissues and diseases.

## Results

### Multiple instance learning for spatial transcriptomics

Our primary approach is to adapt MIL models originally developed for H&E histology, such as **attention-based Multiple Instance Learning (abMIL)** (Ilse et al. 2018) **and Chowder** (Saillard et al. 2020), to SpT data by treating each SpT spot as an instance (Figure 1). Each SpT spot captures the transcriptomic activity of a local region (e.g., in our work, 55 μm diameter with the 10x Visium SD technology), typically heterogeneous mixtures of cells, and is represented by a gene expression vector of raw transcript counts (e.g., in our work, 18,085 genes). Therefore, each SpT sample is represented by a matrix of spot-by-gene expression values (e.g., in our work, n=3,727 spots on average). In our methodology, each spatial spot is first embedded into a lower-dimensional space (e.g., PCA-transformed gene expression or precomputed foundation model features), which is not updated during training of the MIL model. These spot representations are then passed through a neural network composed of fully connected layers to refine the representations for the survival task. For abMIL (Figure 1.B), a gated attention mechanism is used to compute an importance weight for each spot, indicating its contribution to the survival signal. The spot-level representations are then aggregated into a fixed-length patient-level vector via a weighted average using the attention scores (see Supplementary Figure S7.C for the aggregation approach corresponding to Chowder, another MIL model adapted to SpT). This vector is finally passed through a prediction head to produce the survival risk output. We also extended the same principle to scRNA-seq, where the instances are the cell gene expression vectors and each scRNA-seq sample is represented by a cell-by-gene matrix.

**Figure 1.**
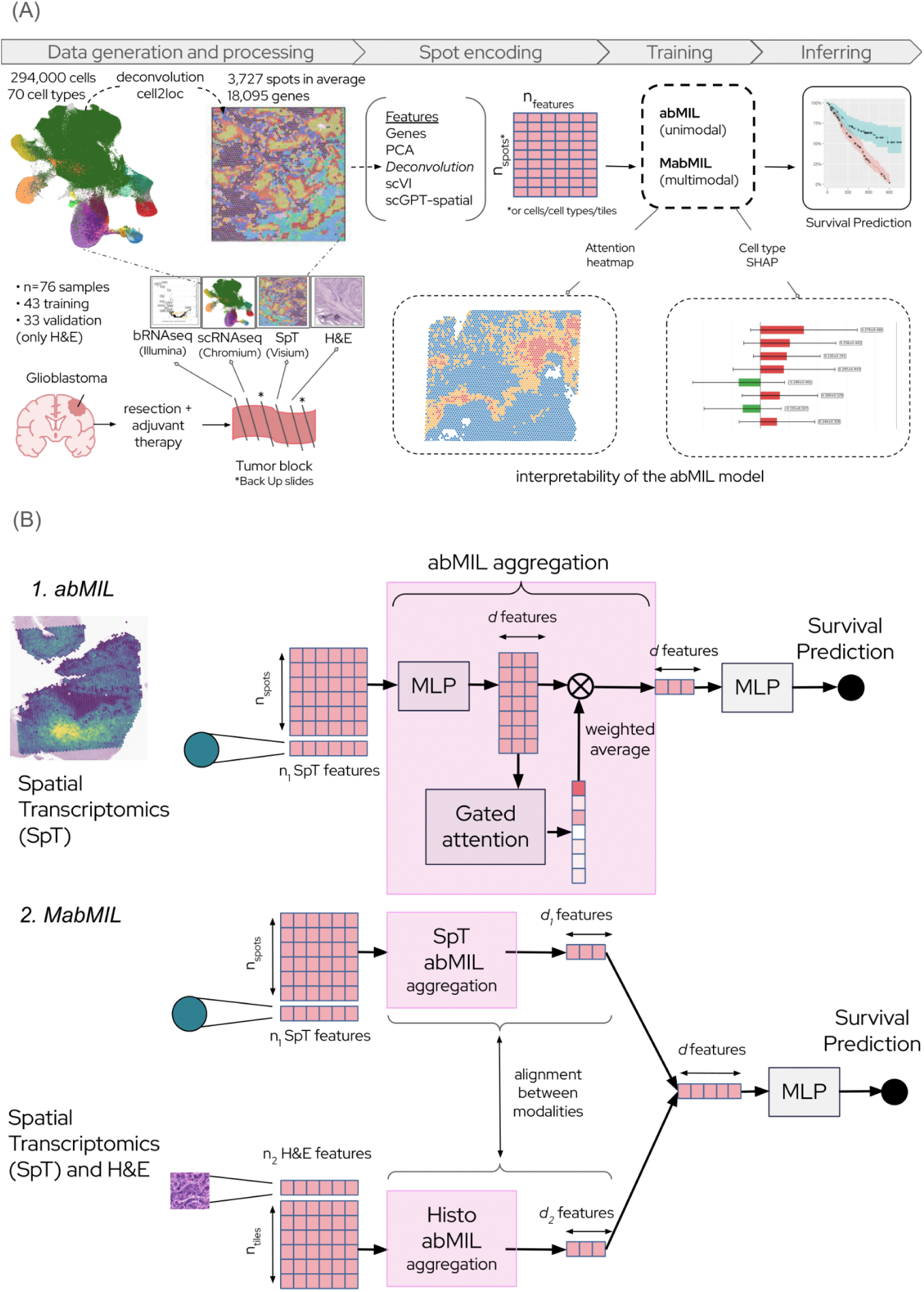
Overview of the workflow and models. (A) Cohort and workflow. The dataset comprised 76 patients, including 43 with matched clinical, bulk RNA-seq, H&E slides, single-cell RNA-seq, and spatial transcriptomics (SpT). Each SpT sample is represented as a spot-by-gene matrix, which can be encoded using various spot representation strategies. These representations are then processed by a Multiple Instance Learning (MIL) model (e.g., abMIL) to predict survival. Fitted models can be interpreted using several approaches; for example, deconvolving each spot vector with reference single-cells enables cell-type–level interpretations via Shapley values. **(B) Attention-based multiple instance learning for spatial transcriptomics.** *1.* abMIL processes SpT spot features through an initial MLP to refine representations, followed by a gated attention network that assigns weights to each spot. A weighted aggregation of spots is then passed through a final MLP to predict survival. *2.* Multimodal abMIL (MabMIL) processes separately SpT and H&E features before aligning them differently depending on the mode: SpT-to-HE distillation or multimodal inference.

Building on this, we implemented multimodal extensions of these MIL architectures for aligned SpT and H&E inputs, enabling joint modeling of transcriptomic and morphological features (Figure 1.B, Supplementary Figure S7.A-F). In the case of abMIL, we introduce the **Multimodal attention-based MIL (MabMIL)** model that uses two parallel abMIL branches to process H&E images and SpT independently, while aligning them through a cross-modal alignment loss. We explore two distinct training strategies for MabMIL (detailed in <u>Results - MabMIL with Spatial transcriptomics to H&E distillation</u>). The first distills prognostic signals from SpT into H&E during training, enabling H&E-only inference. The second strategy leverages spatial alignment between SpT spots and H&E tiles to enable fully multimodal inference.

### SpT outperformed all four other modalities in our overall survival prediction benchmark

We benchmarked the ability of the following five matched data modalities to predict overall survival (OS) (Kaplan and Meier 1958) in 43 GBM patients from the MOSAIC cohort: SpT, H&E, scRNA-seq, bulk RNA-seq, and clinical metadata (Figure 2). OS prediction was evaluated using the concordance index (C-index) (Harrell et al. 1996) in a repeated cross-validation framework, with models and processing tailored to each modality’s structure (as detailed in <u>Methods - Experiment setup</u> and in Supplementary Table S1). Among all modalities and approaches benchmarked, abMIL applied to SpT features achieved the highest predictive performance, with a median C-index of 0.72 and a standard deviation of 0.04. Notably, the reported performance of SpT was significantly higher than bulk, scRNA-seq and H&E features (respective p-values: 0.024, 0.014, 0.0061), but not significantly above clinical data (p-value: 0.32). Clinical variables, such as patient age and MGMT promoter methylation, are currently the strongest individual predictors of survival under SOC treatment (Hegi et al. 2005b; Lacroix et al. 2001). In our benchmark, clinical variables achieved a median C-index of 0.69 (standard deviation: 0.03), making them the second-best performing modality after SpT. Bulk RNA-seq and scRNA-seq features modeled with a Cox proportional hazards model, and H&E features processed with abMIL yielded comparable median C-index values of 0.61, 0.59, 0.58 (standard deviations: 0.05 for all three), respectively (see Supplementary Table S2 for all p-values).

**Figure 2.**
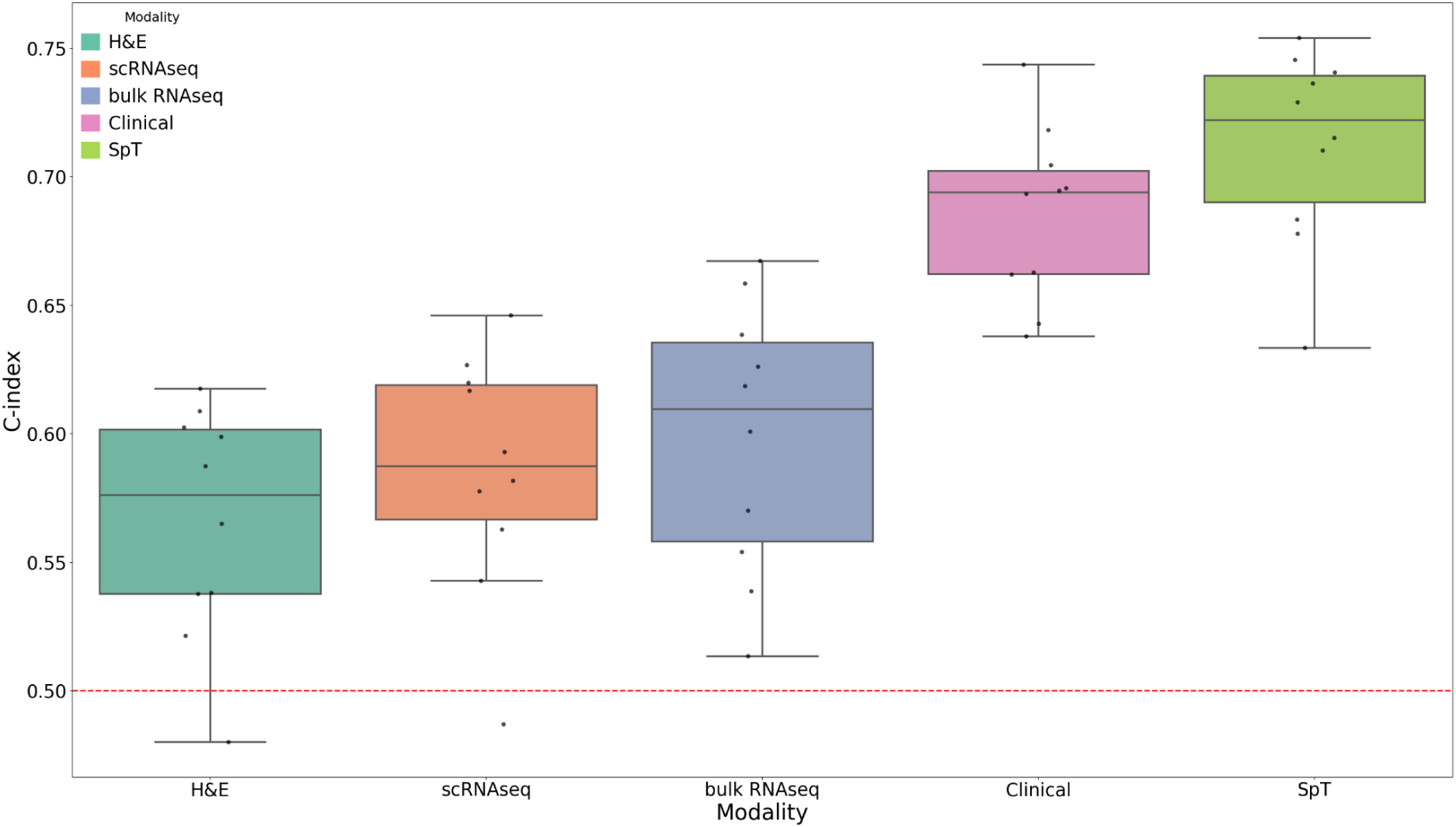
Benchmarking survival models reveals the advantage of abMIL with SpT. For each modality, we evaluated multiple normalization schemes (e.g., raw counts, CPM for SpT), feature transformations (e.g., deconvolution, scGPT-spatial for SpT), and modeling strategies (e.g., abMIL, Chowder for SpT) tailored to its structure. The figure reports the best-performing configuration for each modality (e.g., abMIL for SpT models), ensuring a fair comparison across modalities. From left to right: H&E slides, single-cell RNAseq, bulk RNA-seq, clinical, spatial transcriptomics. Each boxplot represents the distribution of C-index values across repeated 5-fold cross-validation. Each dot corresponds to the mean C-index of one repeat. The red dotted line represents random chance performance.

Interestingly, scRNA-seq reached its best performance (median C-index: 0.59; see above) using a simple pseudobulk approach, where raw counts were averaged across cells for each sample and modeled with a Cox proportional hazards model, rather than a MIL framework (see Supplementary Table S1 for all configurations tried for all modalities). Additionally, we compared pseudobulk-based performances across all molecular modalities (see Methods - Comparison of pseudobulk aggregations). As shown in Supplementary Figure S1, pseudobulks derived from spatial transcriptomics (median C-index: 0.64, standard deviation: 0.03) outperformed pseudobulks derived from scRNA-seq (median C-index: 0.59) or bulk RNA-seq results (median C-index: 0.61), although the performance differences of SpT with scRNA-seq and bulk were not statistically significant here (respective p-values: 0.20 and 0.26). Notably, SpT-based pseudobulk underperformed SpT-based abMIL (median C-index: 0.64 vs 0.72; see Supplementary Table S3 for all p-values). We propose that abMIL can be interpreted as a learned weighted pseudobulk approach, where spot contributions are weighted according to their learned prognostic importance which leads to optimal prognostic accuracy.

### PCA and cell-type deconvolution outperformed other spot representations in SpT MIL models

The effectiveness of MIL-based survival prediction from SpT relies critically on the quality of individual spot representations. Since MIL treats each spot as an instance and aggregates their representations to predict patient-level outcomes, the expressiveness and relevance of the initial spot-level features directly influences model performance. Poorly structured or noisy spot representations can hinder the model’s ability to identify prognostic regions, whereas well-initialized representations can enhance interpretability and downstream prediction. In this section, we therefore systematically evaluated a range of feature extraction strategies for SpT data with the goal of identifying representations that maximize prognostic accuracy in the MIL framework. We evaluated the following transformations within the repeated cross-validation framework for abMIL: Raw counts (no transform, i.e. Identity), Principal Component Analysis (PCA) (Jolliffe and Cadima 2016), Cell-type deconvolution weights per spot (using cell2location (Kleshchevnikov et al. 2022) and paired scRNA-seq), scVI (Lopez et al. 2018) latent representations (fine-tuned per fold), scGPT-spatial representations (Wang et al. 2025) (used in a zero-shot, pre-trained fashion) (see <u>Methods - Foundation models for SpT</u> and Supplementary Table S1). Before going through the representation model, the data were normalized and restricted to a limited number of genes for all methods. More specifically for SpT, we expanded the definition of most variable genes by considering the genes with the highest spatial variability (see <u>Methods - Selection of</u> <u>genes for SpT</u>).

As shown in Figure 3, raw counts performed worst (median C-index: 0.60, standard deviation: 0.02), confirming the necessity of denoising and dimensionality reduction. scVI, despite being fine-tuned for our SpT data, offered limited improvement compared to raw counts (median C-index: 0.63, standard deviation: 0.04) The highest performances came from PCA (median C-index: 0.72, standard deviation: 0.04) and deconvolution features (median C-index: 0.69, standard deviation: 0.04), indicating that compact or biologically informed representations are particularly effective for survival prediction. Interestingly, scGPT-spatial, even without fine-tuning, performed competitively (median C-index: 0.67, standard deviation: 0.04), suggesting that generative transformer models can capture transferable transcriptomic structure (see Supplementary Table S4 for all p-values). The choice of spot representation also impacted runtime at training. Notably, abMIL training with deconvolution features was 2–5x faster than with any other spot representation (see Supplementary Figure S2).

**Figure 3.**
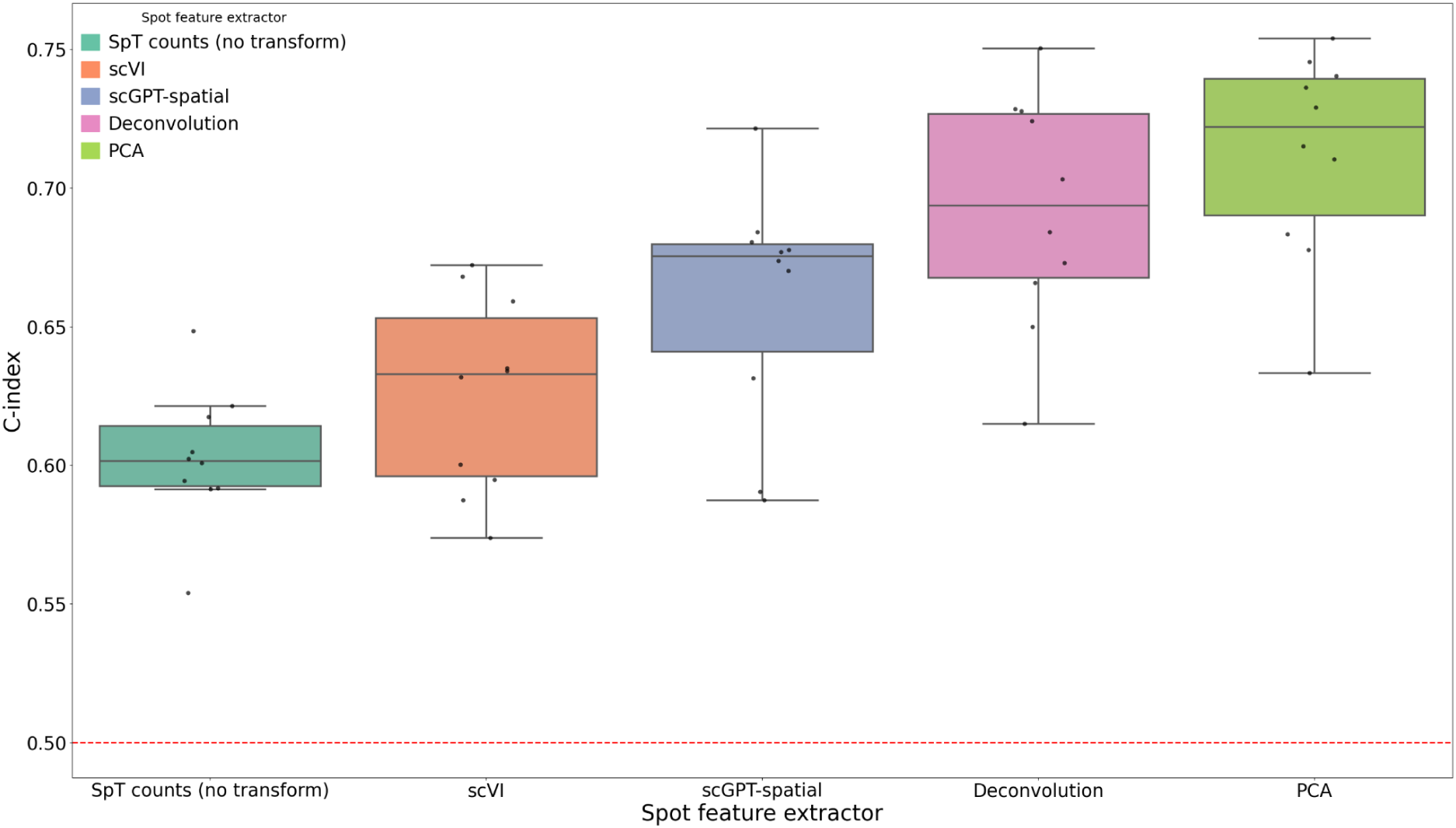
Benchmarking of SpT data transformations for survival prediction. Spatial transcriptomics data were transformed using various strategies, shown left to right: normalized raw counts, scVI, scGPT-spatial (zero-shot), cell-type deconvolution and PCA. Each boxplot represents the distribution of C-index values across repeated 5-fold cross-validation. Each dot corresponds to the mean C-index of one repeat. The red dotted line represents random chance performance.

To better understand the observed differences in performance, we analyzed the latent space structure of each spot representation. As shown in Supplementary Figure S3.A, untransformed SpT counts clustered strongly by sample in the UMAP (McInnes et al. 2020) space, suggesting the presence of substantial batch effects. This was further quantified using the AvgBATCH and AvgBIO metrics introduced in scGPT and scGPT-spatial (Cui, Wang, Maan, and Wang 2023; Wang et al. 2025), where an ideal representation maximizes AvgBATCH (indicating minimal batch-associated variance) and AvgBIO (indicating strong preservation of biological signal). For untransformed SpT counts, these metrics were 0.66 and 0.16, respectively (see Supplementary Table S5). Deconvolution, scGPT-spatial, and scVI representations partially mitigated batch effects (AvgBATCH scores of 0.82, 0.85, and 0.79, respectively) while improving biological signal preservation (AvgBIO scores of 0.26, 0.25, and 0.21, respectively). Among them, deconvolution features explicitly model cell-type proportions within each spot, likely explaining their robustness and interpretability. Interestingly, PCA features, despite achieving the best survival prediction performance, did not exhibit improved structure in this analysis, showing similar metrics to the untransformed SpT counts (AvgBATCH: 0.59; AvgBIO: 0.17). This suggests that existing metrics for evaluating representation quality may not fully capture task-relevant information for clinically important endpoints like survival. We also investigated potential batch effects arising from the sequencing center (“center effect”) or survival type, but these proved minor compared to the dominant sample effect (see Supplementary Figure S3.A-D).

In summary, our results emphasize the importance of transforming spatial transcriptomic data prior to supervised modeling. While generative models like VAEs and Transformers offer principled and increasingly powerful approaches, their success still depends on careful architecture choices, task alignment, and dataset scale. Simpler representations such as PCA or cell-type deconvolution remain strong baselines and in our survival task, it outperformed current state-of-the-art foundation models.

### abMIL interpretability for spatial transcriptomics

abMIL is intrinsically interpretable at the instance level - its attention maps highlight the SpT spots that most influence survival predictions for each sample (see Supplementary Figure S4). However, attention maps do not reveal which cell types are associated with a better or worse prognosis. To bridge this gap, we use cell-type deconvolution features as spot representations, since these are naturally interpretable. This allows abMIL to compute Shapley values directly at the cell-type level (see Supplementary Figure S5). Specifically, we used kernelSHAP (Lundberg and Lee 2017) with abMIL to obtain for each SpT sample the contribution of every cell population to the predicted survival risk (see <u>Methods - Shapley</u> <u>value attribution</u>).

For higher interpretability, we used cell-type deconvolution features consisting of cell counts for 69 cell types as estimated using cell2location (Kleshchevnikov et al. 2022), rather than the 50 used in the SpT benchmark. This finer granularity accounts for tumor cell states that are otherwise ignored. In addition, only the tumoral spots were considered for interpretation; grey matter, white matter and necrosis were excluded by a matter detector (see <u>Methods - Matter detector</u>). The set of cell types spans major malignant, immune and stromal populations (astrocytes, malignant cells, cycling oligodendrocyte precursor cells, neurons, microglia, macrophages, endothelial cells, pericytes, T cells). Incorporating these additional features yielded a median C-index of 0.65 (standard deviation: 0.04; see Supplementary Figure S6).

We quantified feature contributions using KernelSHAP, averaging SHAP values across 20 models (4 repeats of 5-fold cross-validation on 43 GBM patients). Cell types were ranked by their average attribution magnitude in Figure 4. Different malignant subpopulations were linked to either worse or better prognosis. For example, Mes-like and Malignant_SPP1 cells were associated with worse prognosis, whereas OPC-like and AC2-like malignant states were associated with better prognosis. Similarly, among non-malignant populations, Macro_PLIN2 was linked to worse survival, while Pericyte_cycling and cDC1_nos populations were associated with better survival. These associations were consistent across folds.

**Figure 4.**
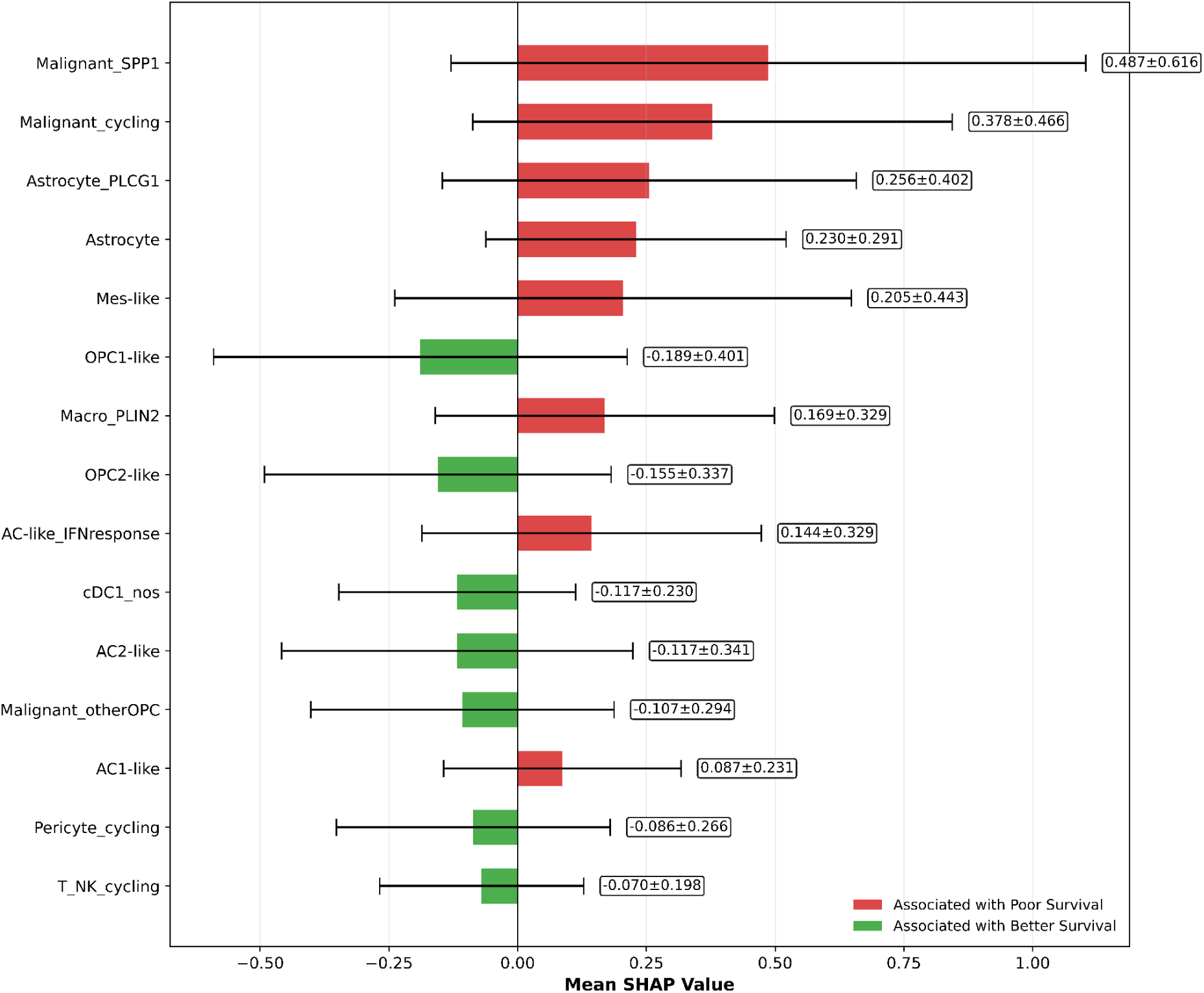
Consensus ranking of cell type contributions to GBM survival prediction using KernelSHAP. This figure summarizes the most reproducible cell type predictors identified across all cross-validation folds. Mean Kernel SHAP values (± standard deviation) were computed across 20 models (4 repeats of 5-fold cross-validation) trained on 43 GBM patients. Cell types are ranked by average attribution magnitude, with red bars indicating association with poorer survival and green bars suggesting protective effects.

### MabMIL with Spatial transcriptomics to H&E distillation

Our multimodal model, MabMIL, can be used in two distinct training and inference paradigms: multimodal inference and SpT-to-H&E distillation (Supplementary Figure S7.A-B).

In the multimodal inference mode, the model processes matched and spatially aligned SpT spot features and H&E tile features through two parallel abMIL branches. Each branch learns to weigh its respective modality via gated attention. The resulting attention-weighted representations are concatenated and used for survival prediction. During training, an additional mean squared error (MSE) loss (Paszke et al. 2019) is applied between the two sets of attention weights to align their attention mechanisms, encouraging the model to focus on concordant prognostic regions across modalities.

In the SpT-to-H&E distillation mode, we leverage the higher prognostic signal from SpT to guide the learning of H&E-based survival prediction. As in the multimodal inference setting, the SpT and H&E features are processed through two parallel abMIL branches. However, only the SpT branch makes the survival prediction, while both branches produce attention-weighted representations. These representations are aligned via a MSE loss, encouraging shared representations across modalities. Notably, in this paradigm, the alignment operates at the aggregated representation level, avoiding the need for explicit one-to-one alignment between individual spots and tiles. During training, the survival loss backpropagates only through the SpT branch, while the alignment loss backpropagates through the H&E branch, establishing a teacher-student dynamic. After training, either branch can be used independently for inference. In our case, we use the H&E branch to enable prediction on unseen histology samples for patients without SpT.

We benchmarked MabMIL by comparing both approaches to abMIL trained on H&E alone, in repeated cross-validation. As illustrated in Figure 5, MabMIL’s multimodal inference mode achieved a median C-index of 0.73 in multimodal inference (standard deviation: 0.05), matching the performance of SpT alone reported previously (median C-index: 0.72). Importantly, MabMIL’s H&E-only inference achieved a higher median C-index (0.59 vs. 0.57) and lower standard deviation (0.03 vs. 0.05) than abMIL trained on H&E alone, although not statistically significant (p-value: 0.32; see Supplementary Table S6 for all p-values).

**Figure 5.**
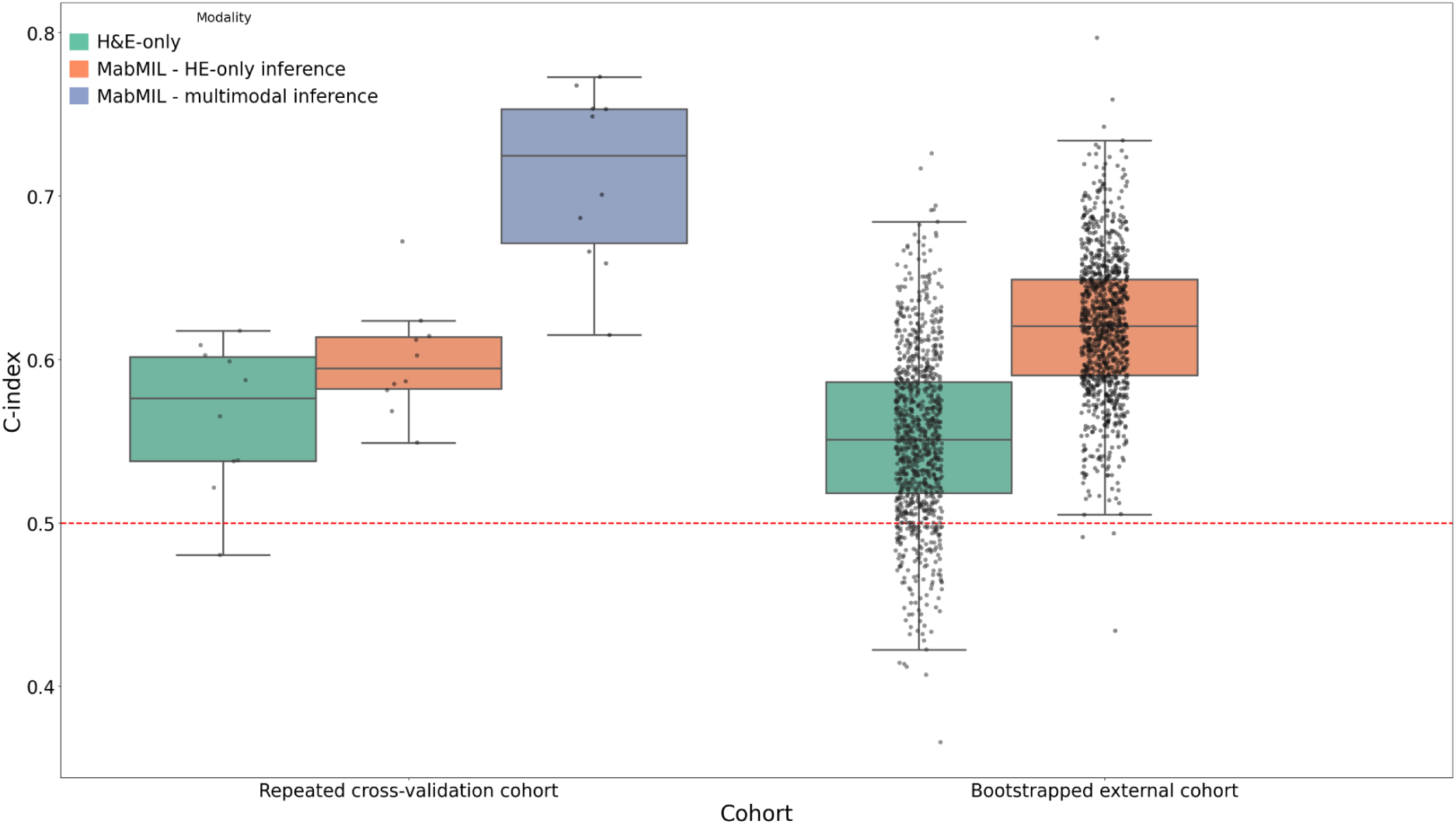
Comparison of MabMIL performances. MabMIL was trained in a multimodal setting (SpT + histology), abMIL (H&E-only) was trained only using H&E. MabMIL (multimodal inference) was evaluated in a multimodal setting (SpT + histology), while MabMIL (H&E-only inference) and abMIL (H&E-only) were evaluated on H&E-data only. The left boxplots represent the distribution of C-index values across repeated 5-fold cross-validation. Each dot corresponds to the mean C-index of one repeat. The right boxplots assess the zero-shot generalization performance of abMIL (H&E-only) and MabMIL (H&E-only inference) on bootstrapped samples from an independent GBM histology cohort, where each point represents a C-index evaluated on a bootstrapped test set. The red dotted line represents random chance performance.

To evaluate generalization of the SpT-to-HE distillation approach, we trained both MabMIL H&E-only inference and abMIL on the full cohort of 43 patients and tested them on an external, unseen cohort of 33 patients with H&E. We assessed zero-shot H&E-only inference by generating 1,000 bootstrap test sets from the 33 external samples. As illustrated in Figure 5, MabMIL again outperformed abMIL and even improved its initial results on the primary cohort, achieving a higher median C-index than abMIL (0.62 [95% CI: 0.54–0.70] vs. 0.55 [95% CI: 0.44–0.66]) and lower standard deviation (0.041 vs. 0.054). Notably, MabMIL was the only model whose performance was significantly higher than the random baseline (C-index = 0.5), underscoring its robustness. These results highlight MabMIL’s ability to distill prognostic signals from SpT into H&E representations, enabling improved H&E-only inference.

## Discussion

This study introduced SpaMIL, a flexible and interpretable MIL framework designed for SpT. Within this framework, we developed abMIL and MabMIL, two attention-based Multiple Instance Learning (MIL) models designed to predict patient-level survival from high-dimensional spatial transcriptomics (SpT) and optionally matched histopathology and scRNA-seq data. We focused specifically on 10x Visium SpT that offers a resolution at spots of diameter 55μm uniformly sampled on a hexagonal grid. By framing SpT modeling as a MIL problem, our models effectively learned prognostic representations from inherently structured data - spots for SpT and tiles for H&E - without requiring uniform input sizes or spatial alignment. To our knowledge, such a level of prognostic accuracy has not been previously reported in the GBM survival prediction literature, where the majority of studies have focused on models leveraging either H&E histology, MRI, clinical metadata or bulk RNA-seq. The modest performance of H&E and bulk RNA-seq was consistent with previous reports in GBM, where prognostic prediction is usually regarded as one of the most challenging tasks (Redlich et al. 2024). Clinical variables, in contrast, are generally considered strong predictors of survival (Chen et al. 2014; Chen, Lu, Wang, et al. 2022; Chen, Lu, Williamson, et al. 2022; Mobadersany et al. 2018), and in our benchmark they indeed achieved high performance. The fact that SpT outperformed clinical features underscores its prognostic power and highlights the potential of spatially resolved transcriptomic profiling to capture biologically and clinically relevant information beyond established covariates. We demonstrated that abMIL significantly outperforms traditional bulk and single-cell RNA-seq approaches in survival prediction. Moreover, naive pseudobulk aggregation of SpT spot-level data outperformed both bulk RNA-Seq and scRNA-seq pseudobulks. These findings highlighted the intrinsic prognostic value of SpT data, suggesting that its added spatial and compositional granularity captures relevant biological signals that are lost in conventional bulk or single-cell resolutions. Each SpT spot contains a mixture of cell types, and the relative composition of these cell types within and across spots provides biologically meaningful information for survival prediction. This may also be reinforced by the fact that, in Visium experiments, pathologists typically select tumor regions of high diagnostic or biological relevance for placement within the capture area, whereas bulk RNA-seq or scRNA-seq often sample from the entire tissue section, potentially diluting signals from the most informative tumor areas.

To our knowledge, this work presented the first adaptation of the MIL framework to SpT. This is likely due to the limited availability of matched SpT data with patient-level survival labels, or more generally to the lack of patient-level matched modalities. Most existing algorithms treat individual spots or cells as independent observations (Wang et al. 2025; Wan et al. 2023; Xu et al. 2024; Wen et al. 2024; Schaar et al. 2024; Bendidi et al. 2024; Hasanaj et al. 2025), which is appropriate for unsupervised tasks like clustering or trajectory inference. However, survival prediction inherently requires learning at the patient level, necessitating models that can aggregate spot-level information while also avoiding overfitting - something that becomes particularly challenging in the absence of sufficiently large patient cohorts. Our MIL framework addressed this challenge directly by treating the patient as the unit of observation and learning to weight intra-sample heterogeneity appropriately.

Our results further show that MabMIL, which integrates SpT with co-registered H&E slides maintained performance comparable to SpT alone in its multimodal inference setting, suggesting that while H&E provides complementary structural and phenotypic context, this additional information may not always boost predictive accuracy beyond what is already captured by local transcriptomic profiles. In its SpT-to-H&E distillation setting, MabMIL managed to distill prognostic signals from SpT into H&E representation. This approach improved the robustness and generalizability of histology-based predictions, as evidenced by the consistent performance gains on an external patient cohort. This has major practical implications: SpT data are scarce and costly, while H&E slides are routinely collected at scale. MabMIL bridges this gap by transferring survival-relevant molecular insights into a modality already ubiquitous in clinical workflows, making spatially-informed survival prediction broadly accessible and clinically scalable.

Our benchmark also explored the impact of different SpT representations on abMIL prognostic accuracy. Despite their promise, existing benchmarks comparing such representation models often rely on limited datasets and only study low-level downstream tasks such as spatial clustering, annotated region classification, and spatial composition estimation (Wang et al. 2025; Wan et al. 2023; Xu et al. 2024; Wen et al. 2024; Schaar et al. 2024). This motivated our own benchmarking study, designed to assess the predictive value of different SpT data transformations for patient-level survival modeling using abMIL. While deep generative models - e.g. scVI (Lopez et al. 2018) and SpT foundation models such as scGPT-spatial (Wang et al. 2025) - offer flexible, spatially aware representations of SpT data, their performance remained below that of simpler representations such as PCA or cell-type deconvolution for survival prediction in GBM. These findings are in line with recent work on foundation models for outcome-related tasks: transcriptomics foundation models underperform methods based on PCA or prior knowledge for survival prediction (Gross et al. 2024) or perturbation response prediction (Bendidi et al. 2024; Hasanaj et al. 2025; Ahlmann-Eltze et al. 2024), and scaling the size of H&E foundation models and their pre-training datasets was reported to not yield consistent gains in survival prediction performance (Filiot et al. 2024). This suggests that, despite their architectural sophistication, current state-of-the-art foundation models may still be limited by training regimes, lack of strong supervision, or sensitivity to technical variation when applied to outcome-related downstream tasks such as survival prediction. However, our results demonstrated that the choice of spot representation has a substantial impact on survival prediction performance, with transformed representations significantly outperforming raw counts. This highlights the importance of evaluating prognostic utility in future foundation model benchmarks, beyond traditional reconstruction or clustering metrics.

By design, our MIL framework embraces the heterogeneity of spatial spots. This is especially crucial in complex malignancies like GBM, where intra-tumoral diversity in cell states and microenvironments may hold prognostic relevance (Neftel et al. 2019). The use of attention weights and *post hoc* cell-type deconvolution based Shapley analysis in abMIL allows for systematic identification of cell types driving patient outcomes, thereby offering an interpretable bridge between computational models and biological insights. For malignant subpopulations, the association of Mes-like states with worse prognosis and OPC-like or AC2-like states with better survival is consistent with previous literature (Neftel et al. 2019; Hara et al. 2021; Ruiz-Moreno et al. 2025). Similarly, among non-malignant cells, Pericyte_cycling has been associated with worse patients outcome (Zhou et al. 2017; Zhang et al. 2021; Sena et al. 2018; Oudenaarden et al. 2022) while the presence ofcDC1_nos population was linked to improved survival (Bowman-Kirigin et al. 2023; Böttcher and Reis e Sousa 2018). By contrast, the associations of Malignant SPP1 and non-malignant Macro_PLIN2 with worse survival were only very recently highlighted (Fidon et al. 2025), making them novel findings in GBM. Crucially, our AI-driven pipeline was able to recover them automatically, without requiring extensive pathologist annotations or manual curation, underscoring both its robustness and practical applicability.

While our study is limited by the number of patients with matched SpT, H&E, single-cell RNA-seq, bulk RNA-seq, and clinical variables, it sets a precedent for the utility of SpT in clinically relevant machine learning pipelines. Future work should explore scaling these models to larger cohorts, incorporating additional modalities such as proteomics, and extending our multimodal distillation approach to other tissues, cancer types or disease phenotypes. Notably, our SpT-based models did not incorporate the spatial coordinates of the spots, relying solely on the gene expression vectors aggregated per spot. This finding highlights the intrinsic prognostic value of spatially localized transcriptomic profiles, even without modeling their physical arrangement, and points to the incorporation of spatial topology as a promising direction to further improve performance. As spatial omics technologies become more accessible, methods like abMIL and MabMIL offer a path forward for developing robust, interpretable, and clinically applicable prognostic tools that leverage the full richness of tissue architecture and molecular content.

## Methods

### Input data

#### MOSAIC Cohort Description

The data consisted of 76 samples at baseline of patients diagnosed of Glioblastoma (GBM IDH-wt), which are part of the MOSAIC consortium dataset, a non-interventional clinical trial registered under NCT06625203. For an overview of MOSAIC data flow and architecture, please refer to the Supplementary Figure S1 of the consortium paper (MOSAIC Consortium and Hoffmann 2025). The GBM samples used in our study originate from two centers: n=24 at University Hospital Erlangen (UKER) located in Germany and n=52 at Centre Hospitalier Universitaire Vaudois (CHUV) in Switzerland. To be included in the MOSAIC GBM dataset, samples needed to be GBM as per WHO 2021 (IDH-wt; H3-wt), primary tumor, patients be aged 45 years or more at diagnosis, treated by debulking surgery followed by adjuvant therapy as per standard of care. Whole Exome Sequencing data was available for these samples as part of the MOSAIC dataset. We meticulously screened for and confirmed the absence of somatic and hotpots mutations in the *IDH1*, *IDH2*, *H3F3A*, *H3C2*, and *H3C3* genes. Samples came from FFPE blocks from resected tumors (minimum 12 micrometers depth and minimum 5×5 mm surface of tissue), and should contain 40 to 80% of tumor content. Sample harvest dearchival ranged from 1.19 to 11.1 years. Samples were excluded if coming from stereotaxic biopsies, cytology or cytoblocks, as well as samples from patients ineligible for surgery and/or with supportive care only. Of note, multiple modalities generated for 10 samples are already publicly available as part of the MOSAIC Window initiative. Among the 76 samples, 43 of them were used to generate multi-molecular modalities and H&E data used for the AbMIL and MabMIL training models, whereas 33 samples (n=19 CHUV, n=14 UKER) were used for the external validation using H&E only.

Clinical and histopathological features were collected across both centers including age, gender, omics information, treatment path, medical history, primary tumor location or number of brain tumor sites. The overall mean age across cohorts was 64.09 ± 10.46 years (CHUV: 66.02 ± 10.29 years, UKER: 59.92 ± 9.74 years). Across both cohorts, 42% of patients (n = 32) were female and 58% (n = 44) were male. A total of 22% of patients had a prior history of cancer (CHUV: 29%, UKER: 8%). In the CHUV cohort, 71% (n = 36) patients underwent maximal safe resection and 29% (n=39) had suboptimal resection, whereas in the UKER cohort, 88% (n=21) underwent maximal safe resection and 12% (n=55) suboptimal resection (see Extended Data Table 1).

#### Data Modalities And Molecular Generation

We studied the MOSAIC GBM tumor biology by incorporating five distinct data modalities: i) Comprehensive clinical data were collected through a dedicated eCRF, meticulously curated, ensuring each patient met specific inclusion criteria. To maintain consistency across the two centers during data collection, clinical features were mapped to a common vocabulary. ii) Microscopic images stained with Hematoxylin and Eosin (H&E) were obtained using standard site-specific protocols. iii) Spatial Transcriptomics (ST) data was generated via the Visium Cytassist standard protocol from 10X Genomics (V4). Single-Nuclei RNA transcriptomics (scRNA-seq) utilized probe-based technology from Chromium Flex (10x Genomics), and Bulk RNA-sequencing (bRNAseq) provided whole transcriptome profiling.

Lab workflow is described extensively in (MOSAIC Consortium and Hoffmann 2025), but briefly, FFPE blocks were reviewed by a pathologist for quality and tumor content, with the optimal block selected and marked for sectioning. Six consecutive sections were prepared for each sample, which allowed optimal alignment between modalities for a single patient: H&E for histology and alignment, Visium sections for spatial transcriptomics (Visium V4, 10x Genomics), and back-up sections. From the remaining block, six curls were cut—two each for single-nuclei RNA-seq (Chromium Flex, 10x Genomics) and bulk RNA-seq. Visium libraries were prepared according to CytAssist protocols; scRNA-seq libraries were generated from isolated nuclei, bulk RNA were extracted using Illumina kits (Figure 1A). All libraries were sequenced at partner CROs, and raw generated data underwent standardized QC and processing pipelines per modality before integration into the MOSAIC dataset.

### Data Evaluation And Processing Steps

#### Histopathology and Image Processing

H&E slides were digitized and quality-controlled to ensure optimal tissue integrity and tumor representation. These images served as spatial references for Visium spot alignment and for guiding downstream macrodissection if required.

#### Bulk RNA Sequencing (bRNAseq)

Bulk RNA-seq reads were aligned to the human reference genome (GRCh38) using STAR v2.7.8a. At this level, samples with more than 40% ribosomal RNA are excluded for downstream applications. Raw read matrix counts were then generated taking into account strandedness of libraries using gene positions and strands from Gencode databases (Gencode v37). Normalisations are then processed from this matrix including TPM(Wagner et al. 2012) and DESeq2 median of ratios (Love et al. 2014).

#### Single-Nuclei RNA Sequencing (scRNA-seq)

10x Genomics Cell Ranger v7.1.0 was used to demultiplex reads and generate gene expression matrices for each sample. To pass QC, samples contained at least 500 cells and more than 50% of reads mapping to the probeset. SoupX v1.5.2 (Young and Behjati 2020) was used to estimate the ambient RNA expression from empty droplets and remove cell free RNA contamination in the data, considering the first 50 principal components. Cells exceeding 10% mitochondrial, ribosomal, or hemoglobin gene content, expressing fewer than 200 genes, or expressing more than 6000 genes were excluded during quality control. Following this cell filtration, genes expressed in fewer than three cells across the entire sample were removed to mitigate potential technical artifacts. Subsequently, scDblFinder v1.4.0 (Germain et al. 2022) with default parameters (19 principal components) was employed to identify and remove multiplet cells. Finally, counts were normalized using normalizations including SCTransform, as implemented in Seurat v4.1.1 (Hao et al. 2024). Individual sample single-cell objects are merged at the level of the dataset to perform cell type annotation by using scVI for alignment (Lopez et al. 2018) and scANVI for transfer cell type annotation from a reference annotation dataset. This reference annotation dataset was composed of all MOSAIC GBM samples, with cells scaled and clustered following a Uniform Manifold Approximation and Projection (UMAP), manually annotated using the cellxgene app (Megill et al. 2021) and known biomarkers of major cell types. Within each major cell type, a Seurat clustering further breaks down cells into distinct transcriptomic profiles, where known or new markers are used to annotate the cells. Similarly, a Seurat clustering was performed on the malignant cluster to annotate malignant subtypes. Some of them were strongly correlated with previously published subtypes based on the gene signature (Neftel et al. 2019; Jong et al. 2025).

The signature matrix at level 4, which was used for downstream deconvolution, was generated using the Seurat FindMarkers function (with default parameters) and the kappa function. This involved differential expression analysis (DEA) between sub-populations of cells and minimization of the condition number. The procedure was performed in a two step approach: i) a first DEA was conducted between cell types at level 2 (low granularity) to capture canonical markers of cell types (e.g. immune cells vs malignant cells) followed by ii) a second DEA between cell types within each group at level 2 (e.g. AC1-like vs AC2-like in the malignant cell group). All genes differentially expressed (based on adjusted p-value < 0.05) are then concatenated and a filtering procedure allowed to keep the signature matrix at optimal size (n=70 marker genes at level 2, n=100 marker genes at level 4, for each cell type). The values in the signature matrix represented the level of expression of those genes in each cell type.

#### Spatial Transcriptomic (SpT)

##### Data processing and integration with scRNA-seq

Sequencing data from Visium CytAssist libraries were processed using 10x Genomics Space Ranger to produce spot-level gene expression matrices, aligned to the reference genome (GRCh38), and spatially registered to the high-resolution H&E image. Quality metrics to include samples included UMI counts, read mapping rates, and spot capture efficiency. Single nuclei RNA-seq–derived cell-type signatures were projected onto SpT spots using the cell2location v0.1.3 deconvolution method on raw counts (Kleshchevnikov et al. 2022), enabling spatial localization of annotated cell types from scRNA-seq.

##### Visium data normalisation

Depending on the experiments, the input SpT data can be either raw counts, or normalized with CPM (counts per million) which removes the effect of sequencing depth per spot. For a given spot *s* and gene *g*, the CPM counts *cpm_s,g_* are derived from the raw counts *c_s,g_* by:

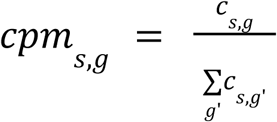

In addition, we optionally apply the x → log(1+x) function to obtain log-CPM counts, which brings the count distribution closer to a Gaussian.

##### Selection of genes for SpT

After the normalisation step, the dimension of the data must be reduced to make it manageable for neural network training. SpT measures the expression of more than 19 thousand genes, which we reduce to less than a thousand, before going through potential transformations like PCA or foundation models. In bulk or single-cell RNAseq, the common way to reduce the dimension is to keep the most variable genes in expression across samples. We propose two different ways to adapt this paradigm to Visium spatial transcriptomics: the first option keeps genes that vary across samples (i), and the second keeps genes that vary spatially within each sample (ii).

i. Variability across samples is estimated by averaging the expression of genes in each sample to obtain a pseudo-bulk sample, and computing the variance of genes across samples.
ii. Alternatively, we compute spatial variability with Moran’s I autocorrelation: this quantity measures how much a gene’s expression in a spot is correlated with its expression in neighboring spots. In a given SpT sample, let *c_i,g_* the counts (raw or normalized) of gene *g* in spot *i*, and *c_g_* the average expression of gene *g* over spots in that sample. Moran’s autocorrelation index *I_g_* in the sample is defined by:

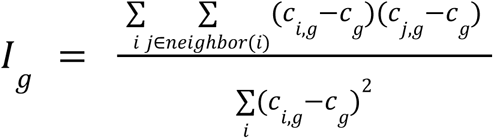

Moran’s I ranges from −1 (perfect anti-correlation) to 1 (perfect correlation), and is close to 0 for random patterns. This quantity is averaged over samples to obtain a ranking of genes by intra-sample, spatial variability.

##### Matter detector

The interpretability framework based on Shapley-value attribution was only done on the spots that were detected as tumor regions by the matter detector; grey matter, white matter and necrosis were excluded. The detector is extensively described in (Fidon et al. 2025), but briefly, we used aligned H&E slides to identify tissue regions with K-means clustering on the H0 tile embeddings of the training cohort. The clusters obtained on the training dataset were annotated by pathologists.

##### Shapley-value attribution

To elucidate the biological drivers behind our MIL survival predictions, we interpreted models trained on deconvolved cell-type and cell-state features (in the case of GBM, tumor-resolved states (Neftel et al. 2019)). We computed feature attributions with both integrated gradients (IG) and KernelSHAP and aggregated results across cross-validation by averaging per-sample attributions over 20 models (4 repeats of 5-fold CV), reporting mean effect ± variability. To ensure spatially coherent biology, we restricted analyses to tumoral spots identified by a tissue clustering algorithm (Fidon et al. 2025), so that cell-type effects reflect only the tumor microenvironment. Concretely, to perform inference with a specific subset of cell types, given the per-spot deconvolution matrix *F* ∈ ℝ^*S*×*C*^ with S the number of spots and C the number of cell types, we construct a binary cell-type mask *m* ∈ {0, 1}^*C*^ with *m*_*c*_ = 1 if and only if the cell type c is in the target subset. We feed the model the masked input *F*′ = *F* ⊙ *m* (mask broadcast across spots), which sets excluded cell-type channels to zero at every spot; i.e.*F*’_*s,c*_ = *F*_*s,c*_ · *m_c_ for s* = 1…*S*, *c* = 1…*C*. Note that no retraining is performed here.

For attribution and “what if” inference, we either sample subgroups *m* and evaluate *f*(*F* ⊙ *m*) (KernelSHAP) or integrate along the path *m*(α) = α · 1, *with* 0 ≤ α ≤ 1 (IG), where 1 is the C-dimensional all-ones vector. The sign and magnitude of the resulting attributions quantify each cell type’s marginal contribution to the prediction of the model with all cell types.

KernelSHAP estimates Shapley values - axiomatically grounded feature attributions - by sampling coalitions of features and fitting a locally weighted linear model around the instance of interest; the Shapley solution satisfies local accuracy, missingness and consistency. It is furthermore model agnostic, capturing non-linear interactions without relying on gradients. In contrast IG integrates gradients along a path from a baseline to the input; while efficient, IG can be sensitive to baseline choice and saturation effects and may under-recover interactions in strongly non-linear regimes. In our setting, IG and KernelSHAP largely agreed on ranked importance and effect direction.

We report KernelSHAP as the primary result because (i) it is model agnostic and therefore robust to architectural details and non-differentiabilities, (ii) it adheres to Shapley axioms ensuring additivity and consistency, and (iii) it exhibited a higher stability across folds in our analysis (smaller variance for top-ranked features), providing a conservative and reproducible summary of prognostic drivers. Positive SHAP values indicate association with poorer survival (risk-increasing), while negative values indicate association with better survival (risk-decreasing).

### Modelisation and Benchmarking

#### Experiment setup

The primary patient-level task benchmarked in this study was the prediction of Overall Survival (OS)(Kaplan and Meier 1958) in individuals diagnosed with GBM. OS was defined as the time from diagnosis (or treatment initiation) to death from any cause. Patients still alive at the time of analysis were treated as censored at their last recorded follow-up. Given the aggressive nature of GBM, only two patients in our dataset were censored, resulting in a nearly complete survival dataset.

Our training and evaluation pipeline was based on repeated cross-validation to ensure robustness and fair comparisons. The 43 patient samples were partitioned with a fixed random seed into five folds for cross-validation. Within each training fold, the most appropriate variables were first filtered, followed by fitting the representation method (except for scGPT-spatial, which was used in a zero-shot setting). The training data were then transformed and used to fit the prediction model. The learned filtering, representation, and prediction steps were subsequently applied to the validation data to generate survival predictions. Each model was evaluated using the concordance index (C-index) (Harrell et al. 1996), which quantifies the agreement between predicted risk scores and actual survival rankings. The C-index scores were then averaged across validation folds. This procedure was repeated with 10 different random seeds, yielding 10 mean C-index values per method. To compare model performance across repeated cross-validation, we used the corrected resampled t-test (Nadeau and Bengio 2003). Unlike a standard t-test, this method accounts for the dependence between folds and repeats, avoiding overly optimistic p-values when evaluating whether one model consistently outperforms another.

For each benchmark, the best-performing configuration was selected through extensive grid search hyperparameter tuning (Bergstra et al., n.d.), ensuring fair comparisons across modalities (see Supplementary Table S1). In particular, we performed more exhaustive searches for scRNA-seq and SpT modalities, as— to the best of our knowledge—such systematic tuning has not been previously reported. For the other modalities, we followed established benchmarking guidelines to constrain the search space. For example, (Gross et al. 2024) have demonstrated that PCA remains a competitive representation method for bulk RNA-seq when coupled with Cox regression (Cox 1972) for survival prediction. Similarly, for histology, we adopted H0 (Saillard et al. 2024) as the feature extractor, which consistently outperforms alternatives in survival benchmarks, coupled with abMIL (Ilse et al. 2018). Clinically relevant variables were selected based on prior studies (Hegi et al. 2005b; Lacroix et al. 2001).

The grid search spanned preprocessing, normalization, feature selection, representation, and survival modeling. For each model, hyperparameters such as the number of selected features, latent dimension size, and predictor settings were tuned. All deep learning models (except scGPT-spatial, which was evaluated in a zero-shot setting) were trained with minibatch gradient descent, with hyperparameters including learning rate, batch size, weight decay, and training epochs selected by cross-validated tuning.

#### Foundation models for SpT

Generative models, particularly Variational Autoencoders (VAEs) (Kingma and Welling 2013) and Transformers (Vaswani et al. 2017), have become foundational architectures in recent advances in spatial transcriptomics. These models learn to represent high-dimensional biological data by capturing its underlying structure in a compressed latent space, enabling reconstruction, simulation, and downstream inference. Their generative capacity allows for denoising, imputation, and synthesis of transcriptomic profiles, making them highly relevant for sparse, noisy, and heterogeneous omics data.

VAEs are probabilistic generative models that assume an underlying latent distribution for each input (e.g., a spot or a cell), typically optimized using a reconstruction loss combined with a Kullback–Leibler divergence term. They model transcript counts using distributions appropriate for RNA-seq data, such as the Negative Binomial or Zero-Inflated Negative Binomial. A prominent example is scVI (Lopez et al. 2018), a VAE originally developed for single-cell transcriptomics, which learns spot- or cell-level latent embeddings that can be used for downstream tasks like clustering or survival prediction. VAEs are often trained in an instance-wise manner (i.e., spot-by-spot or cell-by-cell), with each instance independently encoded and decoded.

Transformer-based generative models approach the problem from a sequence modeling perspective. In spatial transcriptomics, these models treat expression profiles as sequences of tokens - either genes as tokens and cells/spots as sequences (Wang et al. 2025; Schaar et al. 2024; Han et al. 2024), or cells/spots as tokens within a larger tissue context (Wen et al. 2024; 2023). In the former case, a learnable *<CLS>* token summarizes the entire spot or cell and is used as the global representation. In the latter, sequences of spatially adjacent spots can be modeled jointly, optionally incorporating positional embeddings to capture spatial context. These models are trained to predict masked tokens (genes) or reconstruct distributions over them, thereby learning generalizable representations of transcriptomic structure. scGPT (Cui, Wang, Maan, Pang, et al. 2023), and more recently scGPT-spatial (Wang et al. 2025), are key examples of transformer-based generative foundation models now being adapted to spatial omics.

#### Impact of SpT data processing hyperparameters

Supplementary figure S8 shows the impact of pre-processing choices on final model performance: data normalisation, dimensionality reduction method and output dimension.

The results show normalization matters most: CPM negatively affects model performance whatever the gene filtering (raw count median C-index: 0.72, standard deviation 0.04 vs. log-CPM median C-index 0.66, standard deviation 0.02). On raw data, selecting genes at random results in lower C-index than selecting variable genes (random genes median C-index: 0.67, standard deviation 0.04 vs. most variable genes 0.72, standard deviation 0.04), while the median C-indexes between pseudo-bulk and Moran’s I gene selection are close (Moran’s I median C-index 0.70, standard deviation 0.04, pseudobulk variance median C-index 0.72, standard deviation 0.04).

In addition to performing similarly, these two gene selection methods produce very similar gene sets: across folds, the intersection between the most variable genes (pseudo-bulk) and the genes with highest Moran’s I averaged 350 ± 10 genes out of 500, indicating substantial agreement between the two filtering strategies.

#### Comparison of pseudobulk aggregations

To evaluate the utility of MIL-based approaches to predict survival using SpT or scRNA-seq, we compared different pseudobulk strategies across modalities. These included: simple mean aggregation of SpT, simple mean aggregation of single-cell RNA-seq profiles, Bulk RNA-seq, our attention-weighted pseudobulks produced by abMIL. While a simple mean aggregation of SpT equally weighs all spots, this latter representation is enriched in spatially-informed prognostic signals.

As shown in Supplementary Figure S1, pseudobulks derived from single-cell RNA-seq performed worse than bulk RNA-seq but showed comparable variance (median C-indexes: 0.59 vs 0.61, standard deviations: 0.05 for both). Mean aggregation of SpT spots performed better with standard deviation twice as lower (median C-index: 0.64, standard deviation: 0.03). However, the highest performance was achieved by attention-weighted pseudobulks (median C-index: 0.72, standard deviation: 0.04), the aggregation type used in abMIL and MabMIL, which can learn to prioritize prognostically informative regions of the tissue end-to-end while learning to predict survival during training.

The results confirmed our previous findings that spatial transcriptomics provided inherently stronger prognostic signal than single-cell or bulk RNA-seq - even when spot-level data is aggregated using a simple mean. This suggests that SpT captures biologically-relevant spatial information not preserved in bulk or single-cell data that are informative for survival prediction. By assigning higher weight to the most informative regions, abMIL and MabMIL transform the raw data into optimized pseudobulks that outperform all other representations.

#### Implementation details

All models in the following section were implemented and trained using PyTorch (Paszke et al. 2019), unless stated otherwise.

For H&E, the optimal pipeline involved transforming each tile using the H0 model and sampling 4,000 tiles per slide, rather than restricting to only the tiles aligned with SpT. The best prediction model was abMIL, trained with gradient descent for 50 epochs, a learning rate of 0.001, and no weight decay. The embedding multi-layer perceptron (MLP) (Rumelhart et al. 1986) consisted of one hidden layer with 128 nodes and an embedding dimension of 128, while the prediction MLP also comprised a single hidden layer of 128 nodes.

For scRNA-seq, the optimal pipeline first aggregated all cells per patient to create a pseudobulk representation. The 1,000 most variable genes were then selected, and the raw counts were scaled using the median-of-ratios method from PyDESeq2 (Love et al. 2014; Muzellec et al. 2022). A 20-dimensional PCA was fit, and survival was predicted using a Cox model implemented in scikit-learn (Pedregosa et al. 2011) with a penalizer of 0.01 and a L1 ratio of 0.5.

For bulk RNA-seq, the optimal pipeline filtered the TPM (Wagner et al. 2012) normalized data to the 3,000 most variable genes, before applying a log min-max scaling. A 5-dimensional PCA was fit, and survival was predicted using a Cox model with a penalizer of 0.001 and a L1 ratio of 0.5.

For clinical data, the columns kept for prediction were the administrative gender, age at diagnosis, MGMT promoter methylation status and surgery type. Within each fold, missing values were replaced by the most frequent one (categorical) or the median one (numerical), the binary categories were converted to integers, and the numerical values were transformed following a standard scaling. Survival was predicted using a Cox model with a penalizer of 0.1 and a L1 ratio of 0.1.

For SpT, the optimal pipeline selected the 500 most variable genes according to the pseudobulk approach (detailed in <u>Methods - Selection of genes for SpT</u>), before transforming the raw counts using PCA to a dimension of 500 features (thus effectively rotating the raw counts). The best prediction model was abMIL, trained with gradient descent for 30 epochs, a learning rate of 0.0003, and no weight decay. The embedding MLP consisted of no hidden layer and an embedding dimension of 128, while the prediction MLP comprised two hidden layers of 128 and 64 nodes.

For the interpretation of SpT deconvolution features, the counts were transformed to 69 deconvolution weights by using scRNA-seq-derived profiles and Cell2Location (as described in <u>Methods - Data processing and integration with scRNA-seq</u>). The best prediction model was abMIL, trained with mini-batch gradient descent for 50 epochs, a learning rate of 0.001, a weight decay of 0.0001 and a batch size of 32. The embedding MLP consisted of no hidden layer and an output dimension of 256, while the prediction MLP comprised two hidden layers of 128 and 64 nodes each. After training, only the tumoral spots were kept for interpretability by KernelSHAP.

For multimodal inference models described in Supplementary Figures S7.A-B, the optimal pipeline was obtained with MabMIL multimodal inference mode. The model uses aligned SpT and H&E features, thus being able to compute a MSE loss between both sets of weights. It selected the 500 most spatially variable SpT genes according to the Moran’s approach (detailed in <u>Methods - Selection of genes for SpT</u>), before transforming the raw counts using PCA to a dimension of 500 features (thus effectively rotating the raw counts). The model was trained with gradient descent for 30 epochs, a learning rate of 0.0003, and a weight decay of 0.001. The embedding MLPs of both the SpT and H&E branches consisted of no hidden layer and an embedding dimension of 256 before concatenation, while the prediction MLP comprised two hidden layers of 128 and 64 nodes.

For MabMIL’s SpT-to-H&E distillation mode, the optimal pipeline uses SpT features and H&E features, the latter being the whole set of H&E tiles instead of only the aligned ones. It selected the 500 most spatially variable SpT genes according to the pseudobulk approach (detailed in <u>Methods - Selection of genes for SpT</u>), before transforming the raw counts using PCA to a dimension of 256 features. The model was trained with gradient descent for 30 epochs, a learning rate of 0.001, and no weight decay. The embedding MLP of the SpT branch consisted of no hidden layer, while the embedding MLP of the H&E branch consisted of one hidden layer of 256 nodes, both mapping to a dimension of 128. The prediction MLP comprised two hidden layers of 128 and 64 nodes.

## Supplementary

### Supplementary Figures

**Supplementary Figure S1.**
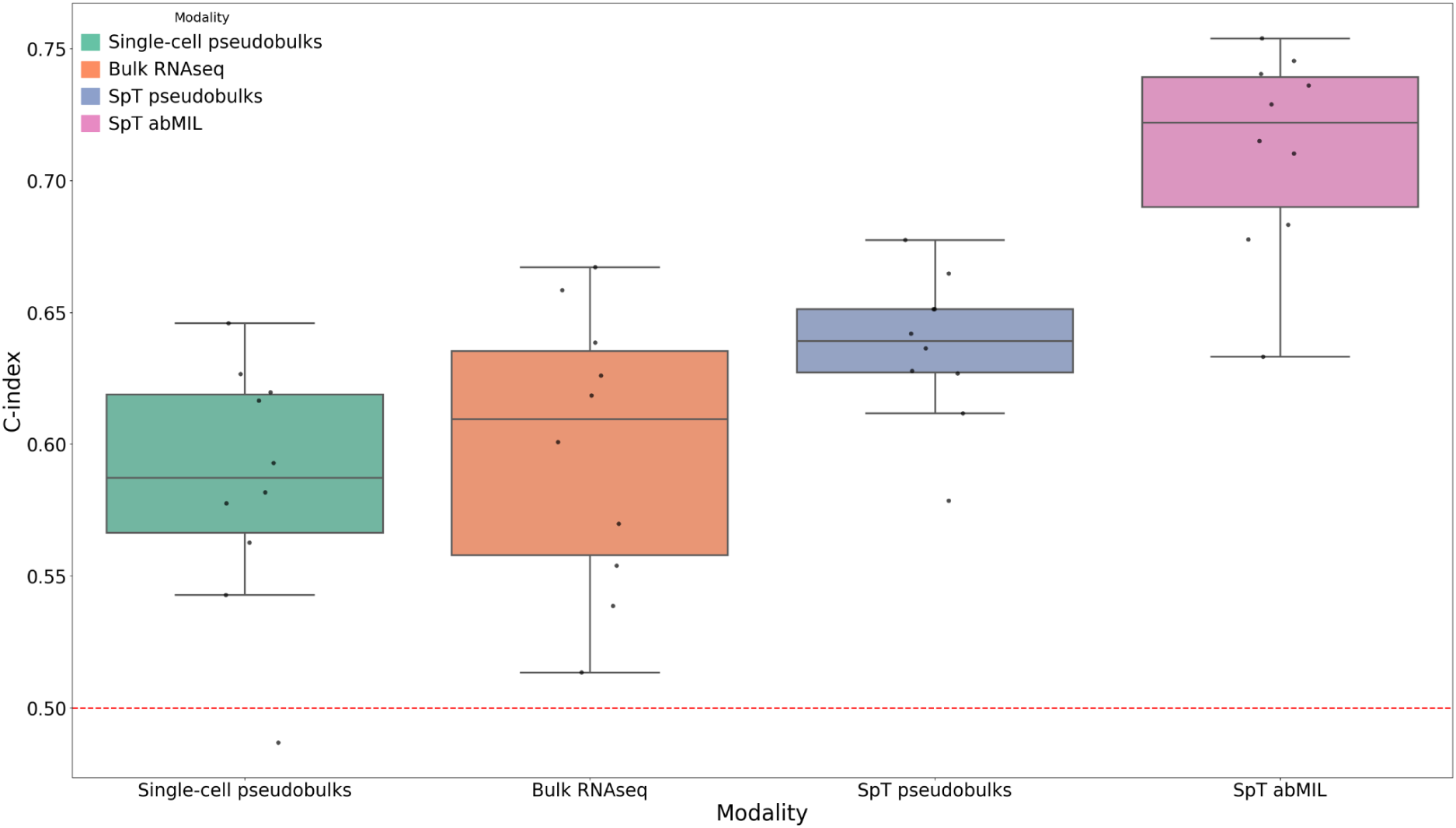
Benchmarking of bulk and pseudobulk aggregation strategies for survival prediction. Various pseudobulk approaches were evaluated to approximate bulk RNA-seq. From left to right: equally weighted pseudobulk profiles derived from single-cell RNA-seq, true bulk RNA-seq, equally weighted pseudobulk profiles derived from spatial transcriptomics, and abMIL attention-weighted pseudobulk profiles from spatial transcriptomics. Each boxplot represents the distribution of C-index values across repeated 5-fold cross-validation. Each dot corresponds to the mean C-index of one repeat. The red dotted line represents random chance performance.

**Supplementary Figure S2.**
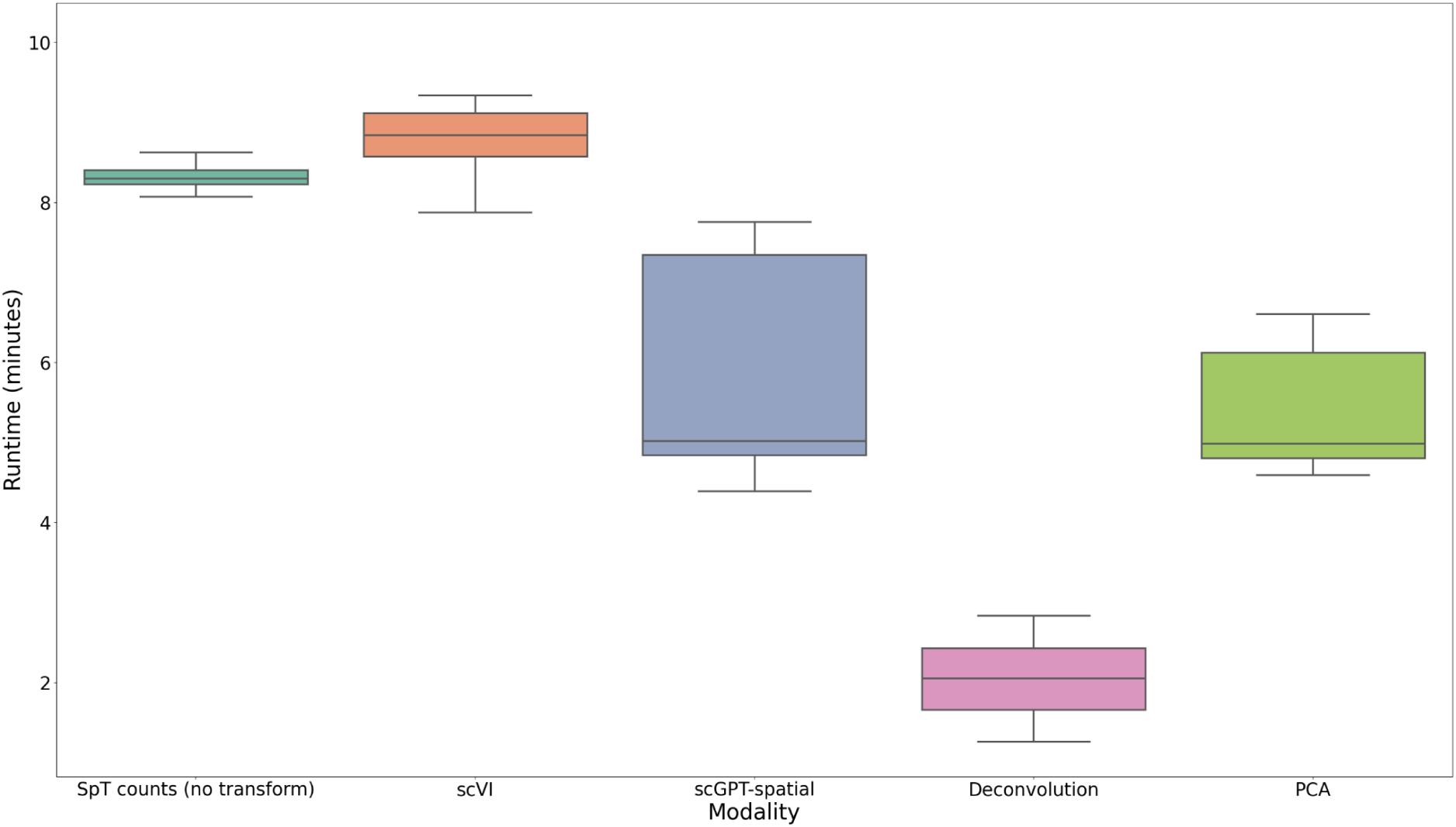
Comparison of abMIL’s training runtime. The runtime of abMIL’s training is evaluated across 10 repeats of 5 fold cross-validation, in minutes. From left to right, abMIL is trained using features from: SpT counts (no transform), scVI, scGPT-spatial, Deconvolution, PCA.

**Supplementary Figure S3.**
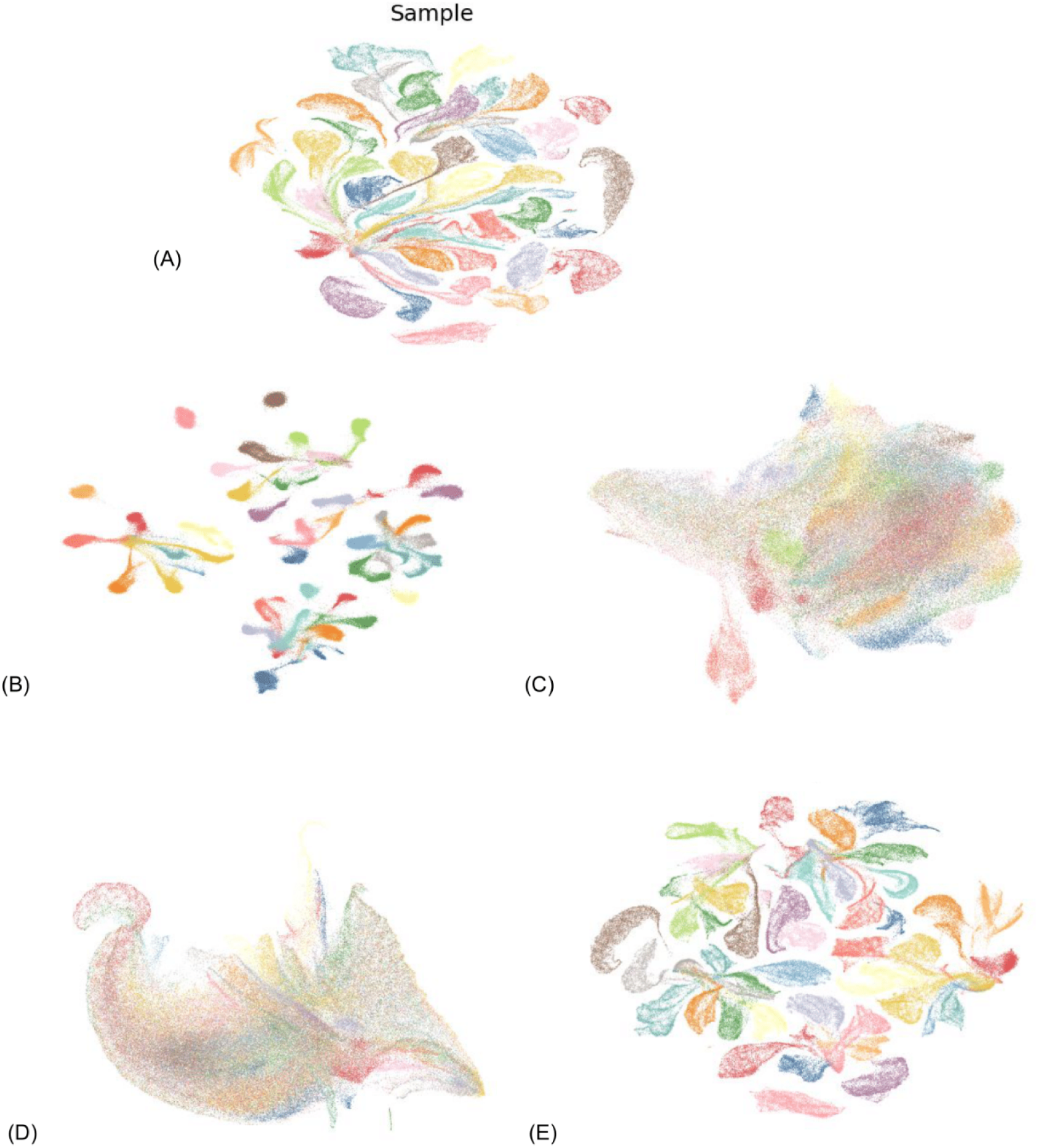
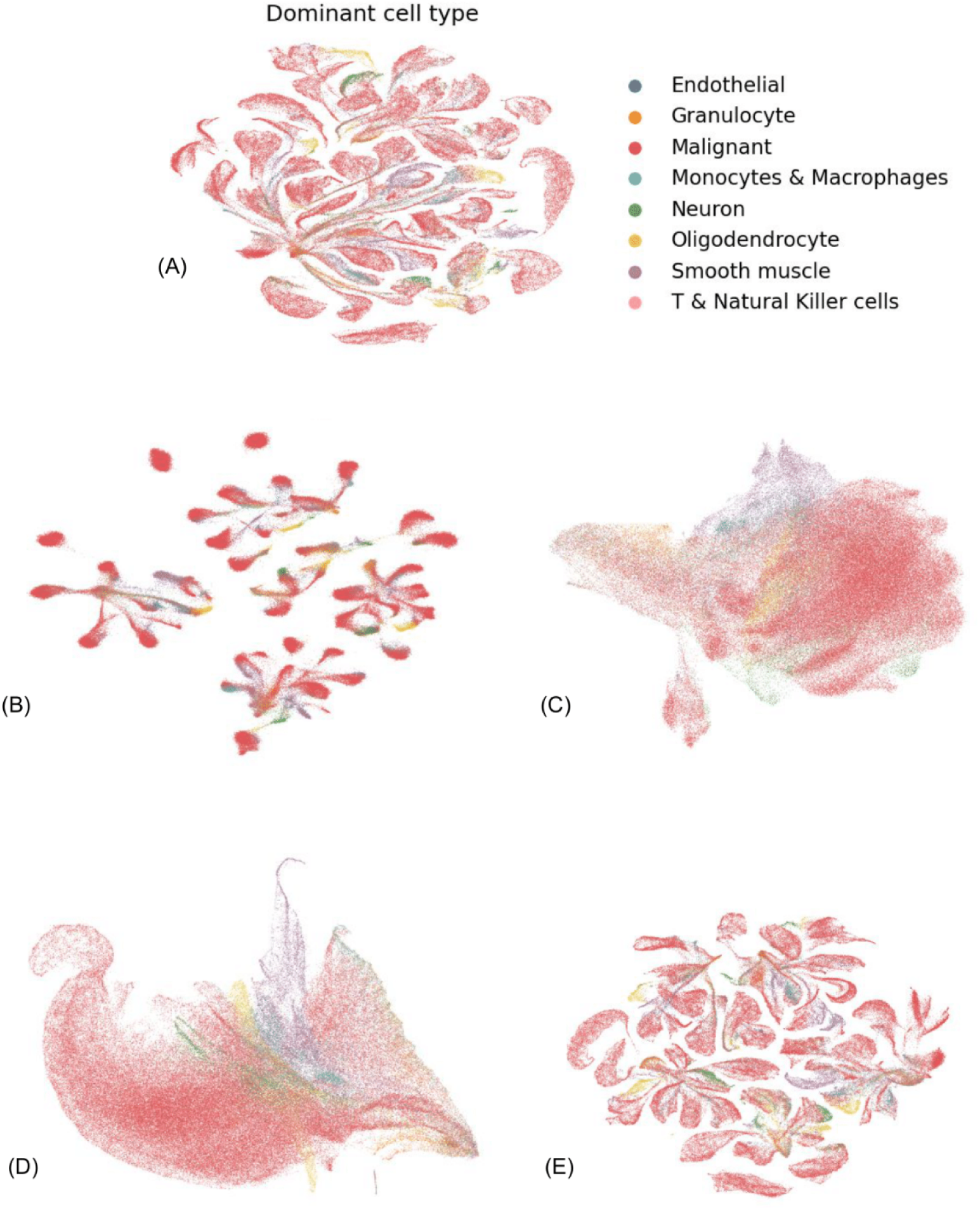
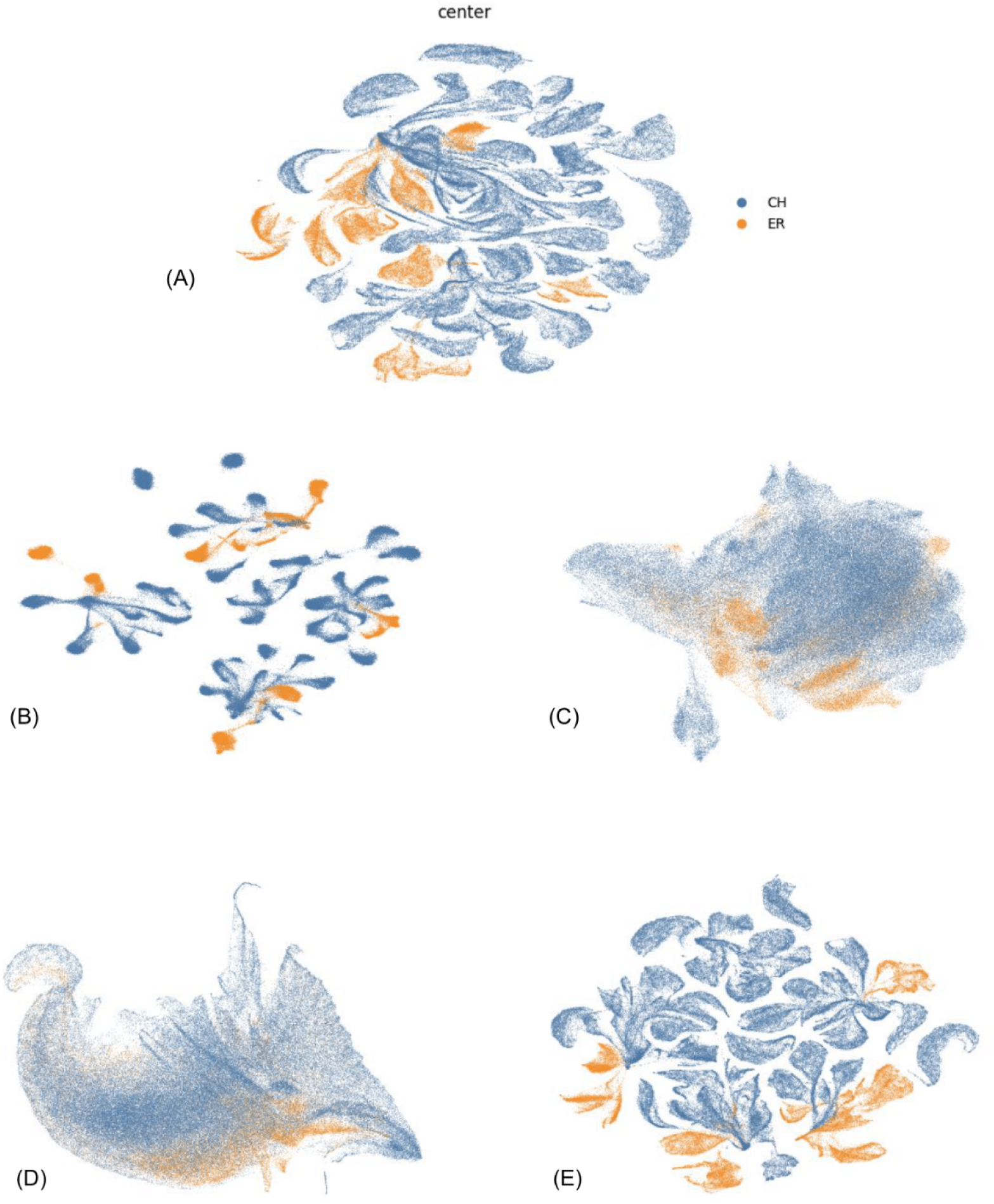
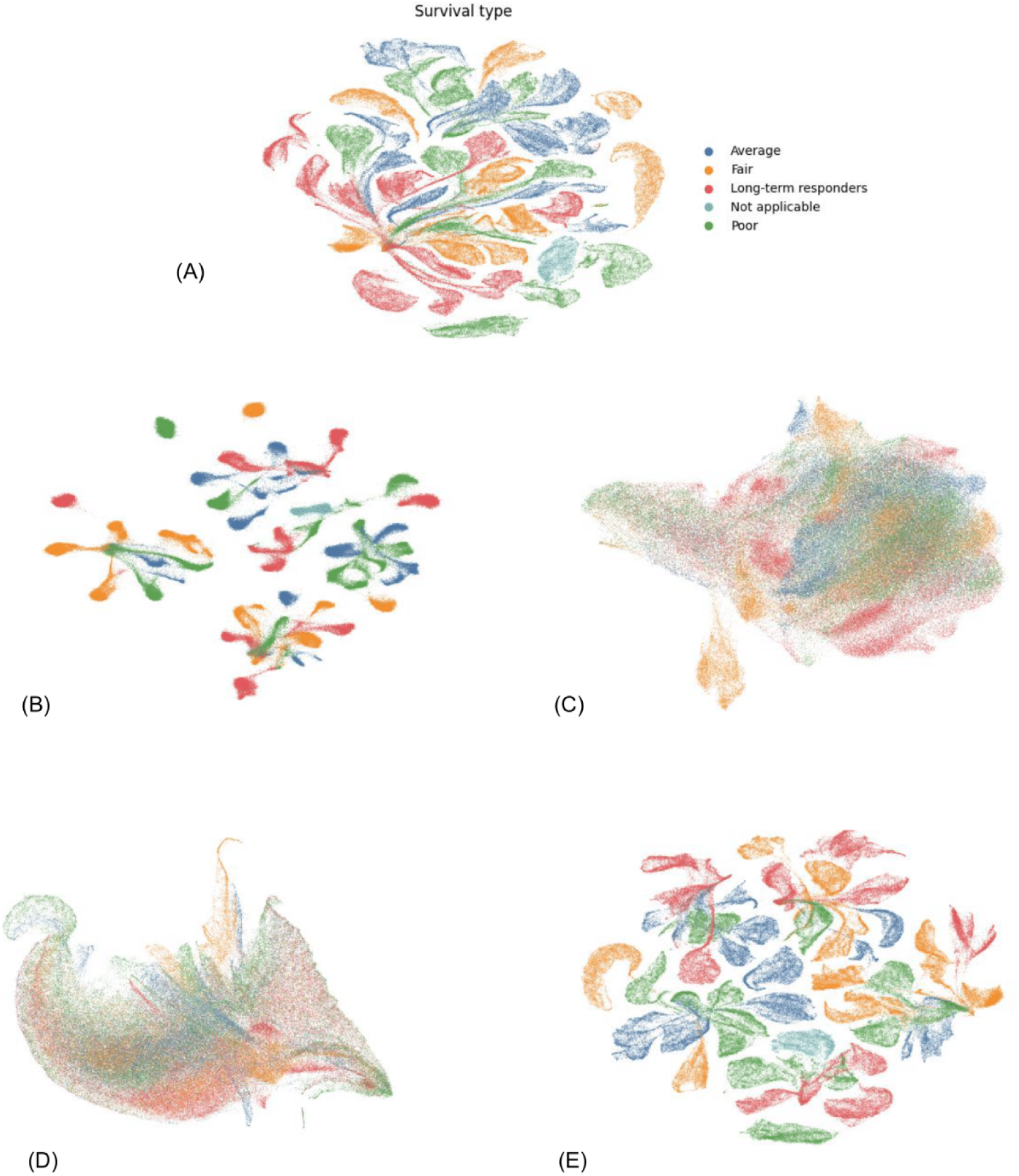
A UMAP representation of the SpT spot counts, colored by sample. Each spot is coloured by the sample it originates from, under different representation algorithms. (A) Counts, no transform. (B) scVI. (C) scGPT-spatial. (D) Deconvolution. (E) PCA. **B UMAP representation of the SpT spot counts, colored by dominant cell type**. Each spot is coloured by the most frequent cell type in that spot, under different representation algorithms. (A) Counts, no transform. (B) scVI. (C) scGPT-spatial. (D) Deconvolution. (E) PCA. **C UMAP representation of the SpT spot counts, colored by center**. Each spot is coloured by the center the sample originates from, under different representation algorithms. (A) Counts, no transform. (B) scVI. (C) scGPT-spatial. (D) Deconvolution. (E) PCA. **D UMAP representation of the SpT spot counts, colored by survival type**. Each spot is coloured by the survival type of the patient. A patient was decided of poor survival for an overall survival between 0 to 0.75 years; of average survival for an overall survival between 0.75 to 1.5 years; of fair survival between 1.5 to 3 years; of long-term survival for more than 3 years; not applicable if censorship occurred for less than 3 years. (A) Counts, no transform. (B) scVI. (C) scGPT-spatial. (D) Deconvolution. (E) PCA.

**Supplementary Figure S4.**
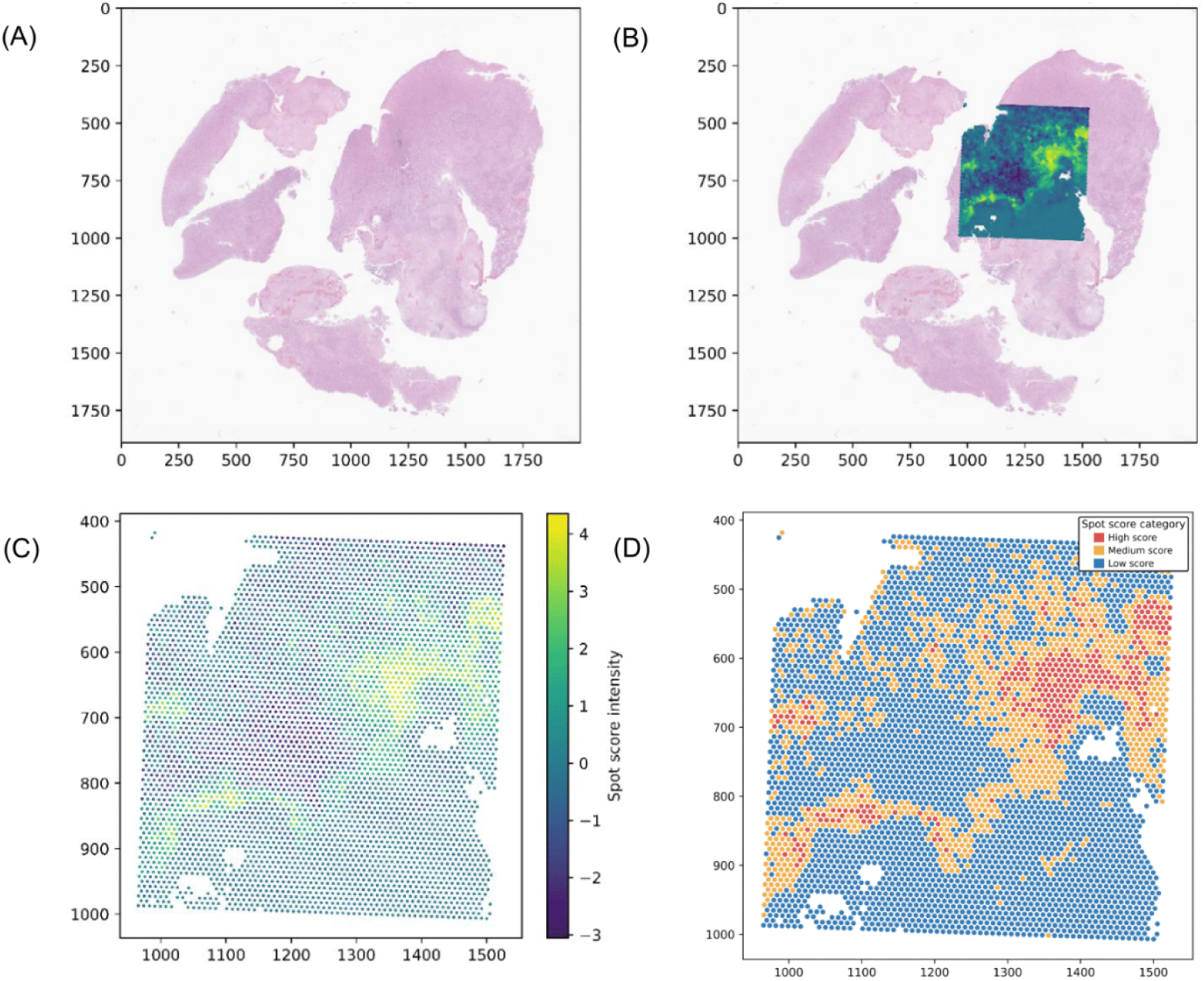
abMIL attention weights create spatial heatmaps. Each tile is weighted by abMIL according to its prognostic importance. (A) A sample’s histology slide. (B) The aligned spot heatmap on the slide, created from the trained abMIL attention weights. (C) The attention heatmap with the spot score intensity. (D) Division of the spot score intensity between high, medium and low scores.

**Supplementary Figure S5.**
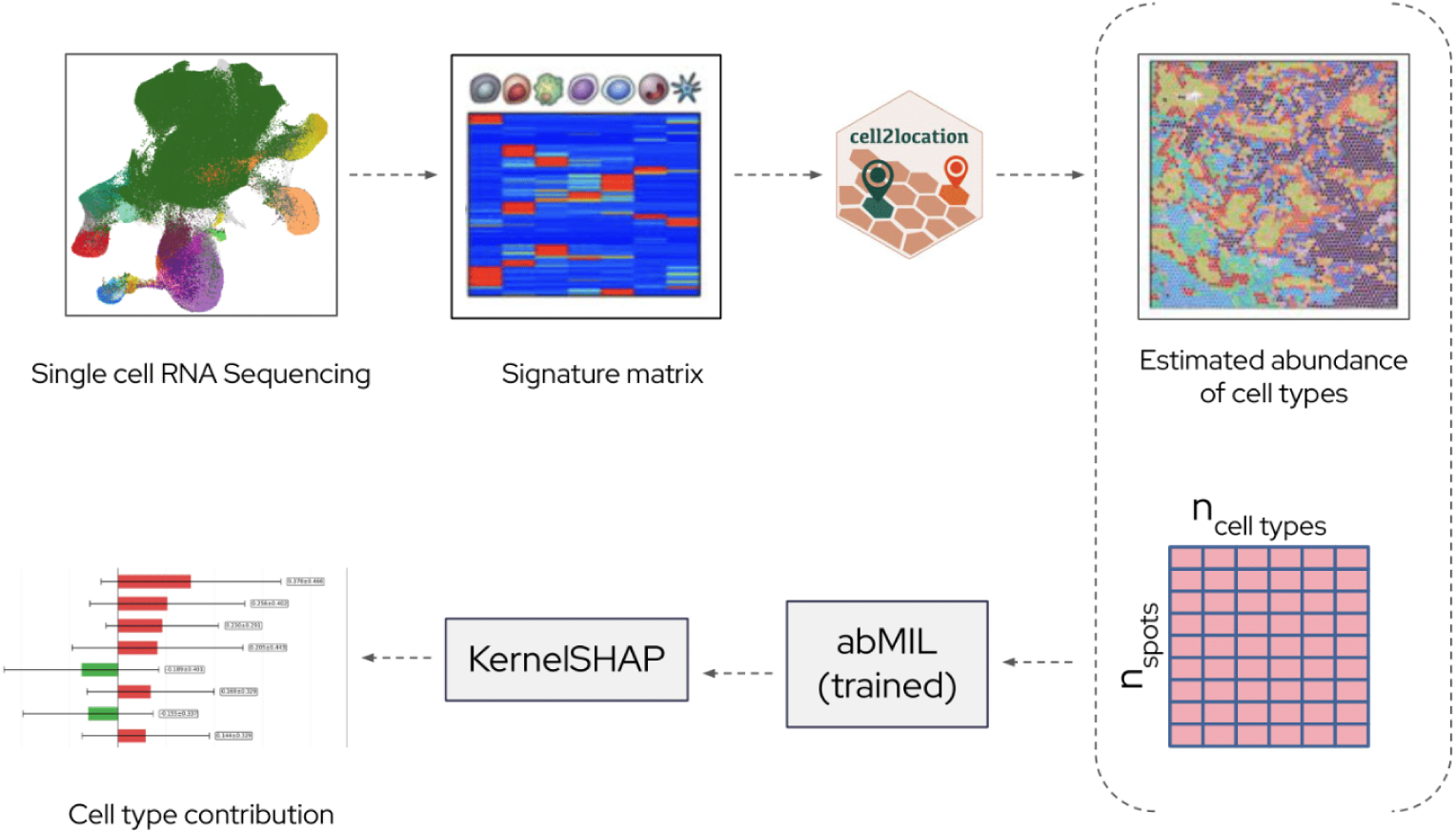
From single-cell references to interpretable spatial predictions. Workflow overview. scRNA-seq data are used to build a cell-type signature matrix, which is provided to cell2location to deconvolve spatial transcriptomics spots and estimate spatial abundance of each cell type. The output is an n_spots × n_cell-types matrix of cell-type abundances (right). These features are then modeled with an attention-based multiple instance learning network (abMIL) to predict section- or region-level outcomes. Model attribution with KernelSHAP quantifies the contribution of each cell type to the prediction, yielding interpretable per–cell-type effect sizes (bar plot). Dashed arrows indicate data flow between steps.

**Supplementary Figure S6.**
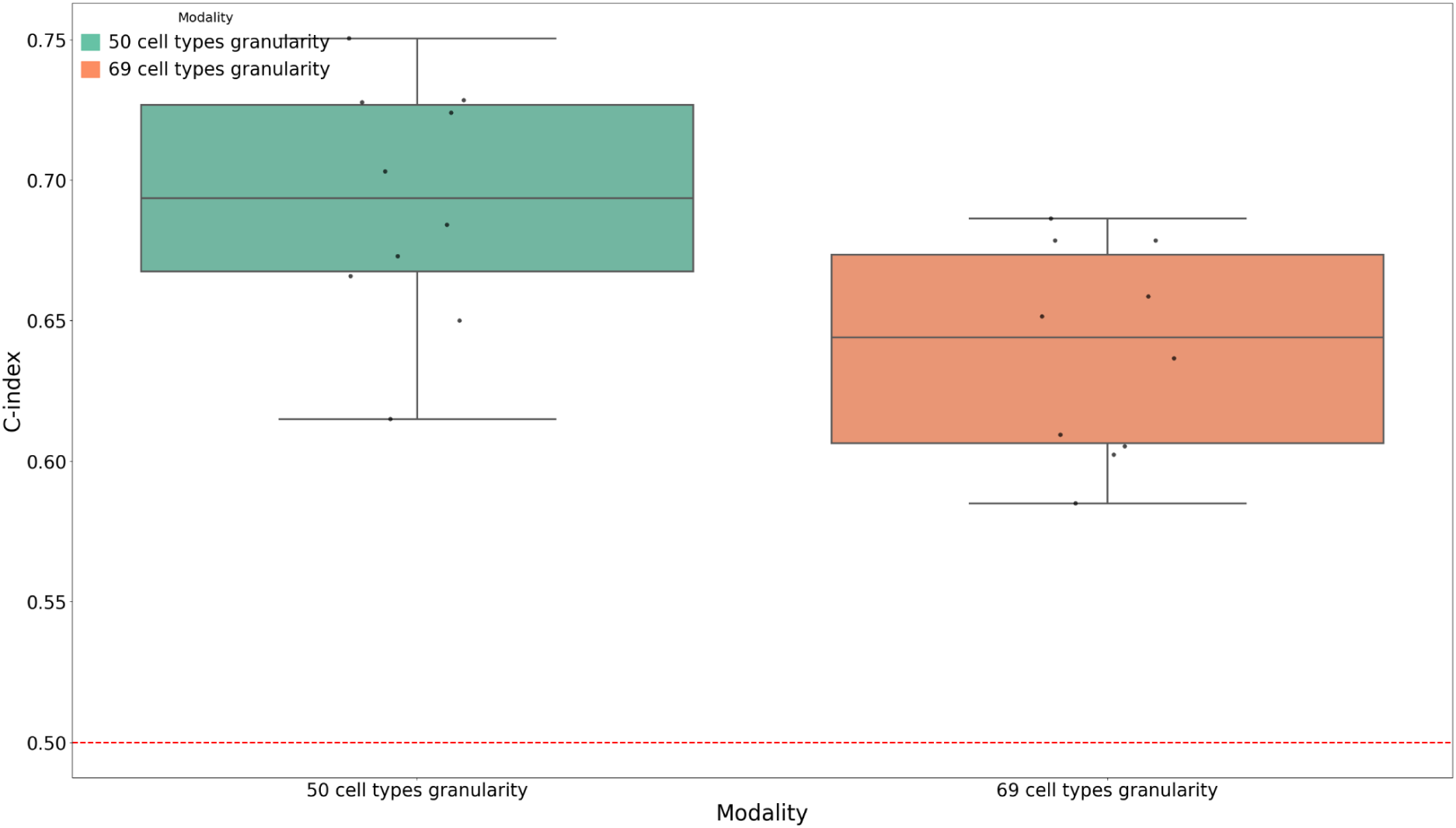
Deconvolution inference for interpretability. The interpretability of the model was done using the highest deconvolution granularity, which includes tumoral subtypes. For this granularity, the inference was only performed on the tumoral spots. The red dotted line represents random chance performance.

**Supplementary Figure S7.**
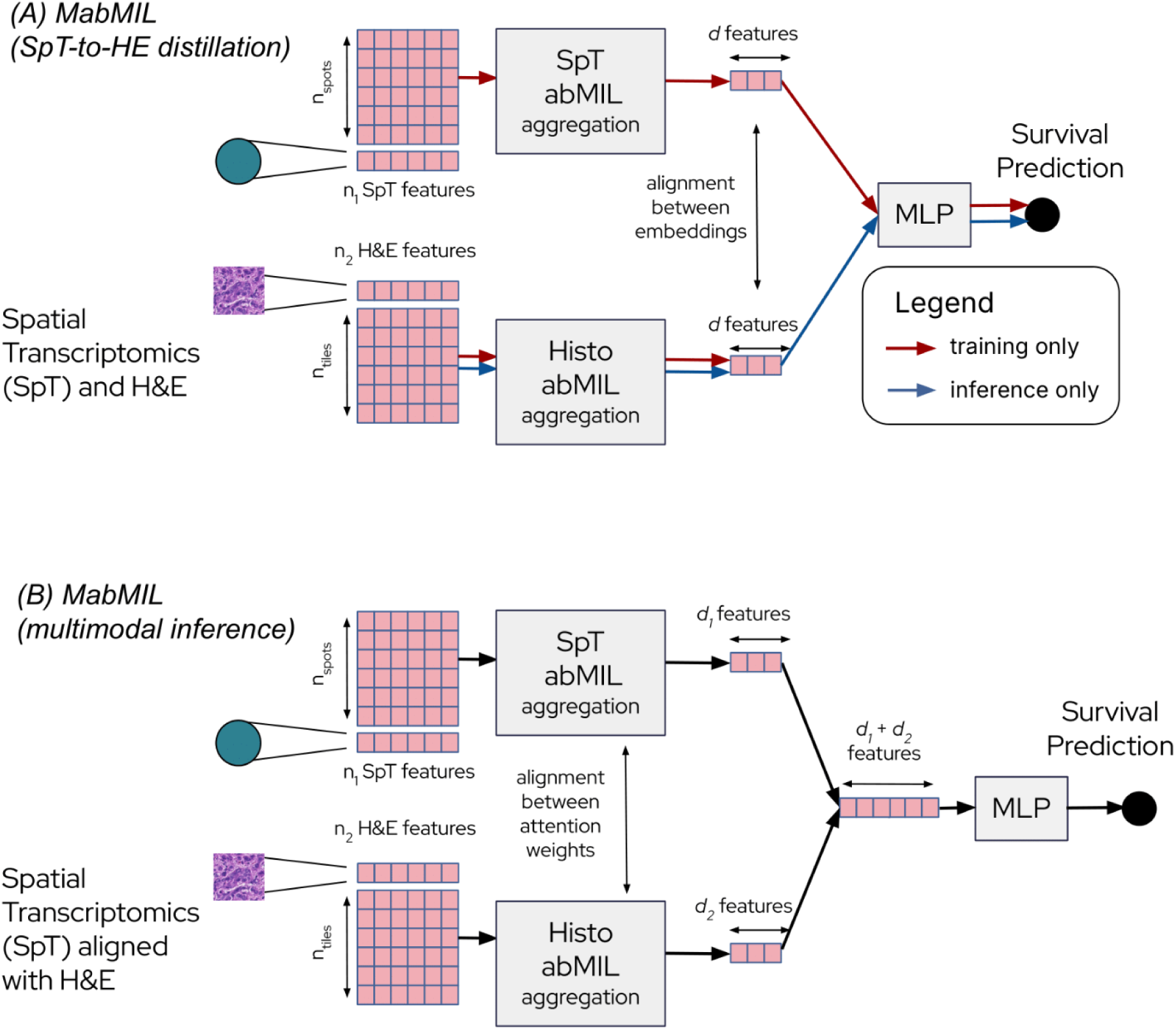
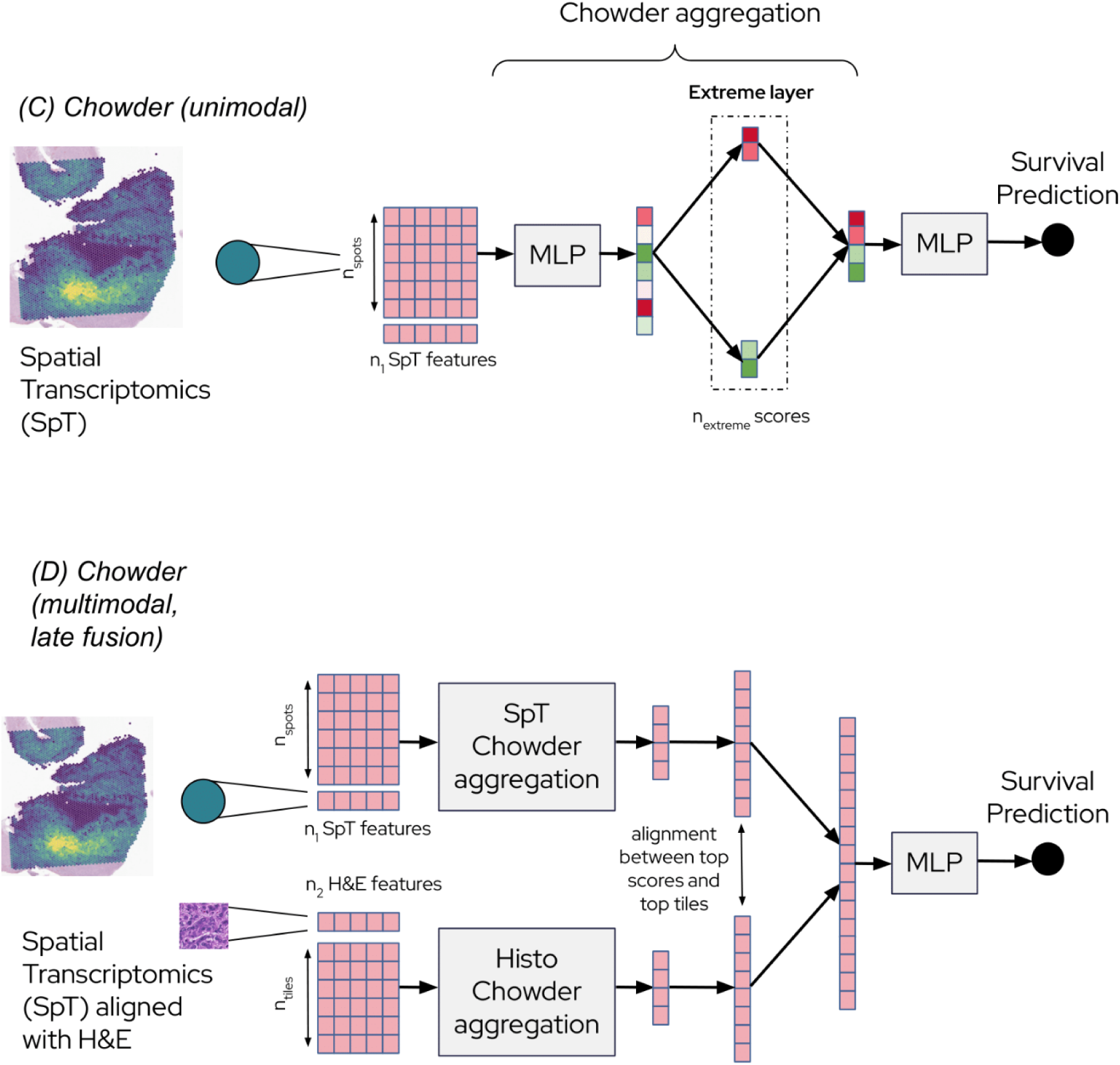
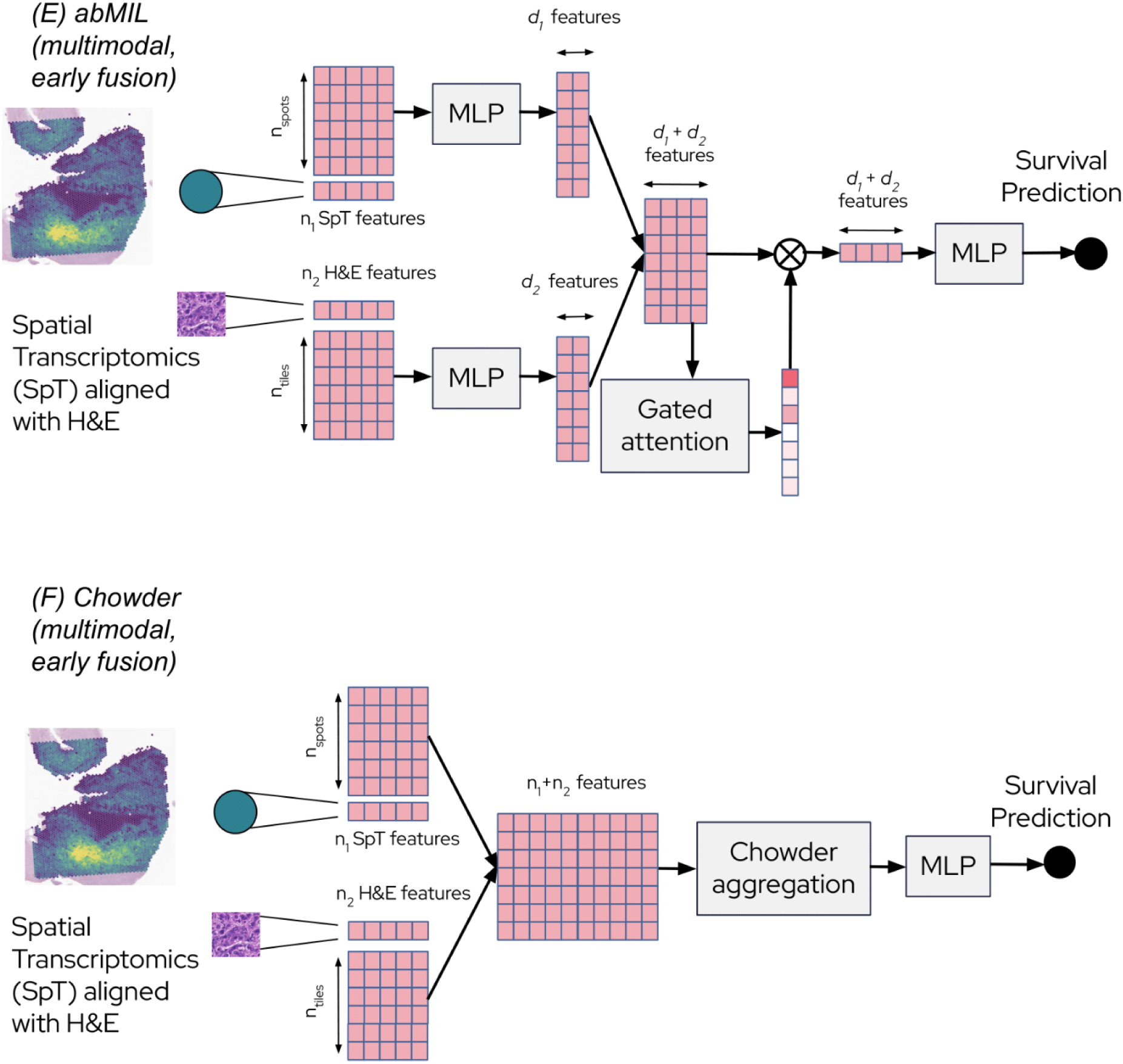
Models schematics. **(A) MabMIL** (SpT-to-HE distillation) workflow. Two abMIL branches are trained in parallel: one processes histology-tile features, the other SpT-spot features. Only one branch outputs the survival prediction (SpT here), but both branches are trained to learn similar attention-weighted representations through a consistency loss. During inference, the model can make predictions using only a single modality (histology here). **(B) MabMIL** (multimodal inference) workflow. Histology and SpT features are again processed by separate abMIL branches, but their attention-weighted embeddings are concatenated and jointly used for survival prediction. When the two modalities are spatially aligned, an additional loss promotes alignment between high-attention histology tiles and matching high-attention SpT spots. **(C) Chowder** ( unimodal) workflow. Each spot is scored through a shared MLP. Only the extreme scores are kept and concatenated to go through a predictive MLP, which outputs survival prediction. **(D) Chowder** (multimodal, late fusion) workflow. Histology and SpT features are processed by separate Chowder branches, their extreme scores are concatenated and jointly used for survival prediction. When the two modalities are spatially aligned, an additional loss promotes alignment between high-scoring histology tiles and matching high-scoring SpT spots. **(E)** abMIL (multimodal, early fusion) workflow. After an initial MLP refining the representation of the SpT and H&E data, each spot and tiles are concatenated and go through the gated attention mechanism as one entity, before their aggregation goes through the predictive MLP to output survival prediction. **(F) Chowder** (multimodal, early fusion) workflow. Each spot and tiles are concatenated and scored as one entity, before their extreme scores go through the predictive MLP to output survival predictions.

**Supplementary Figure S8.**
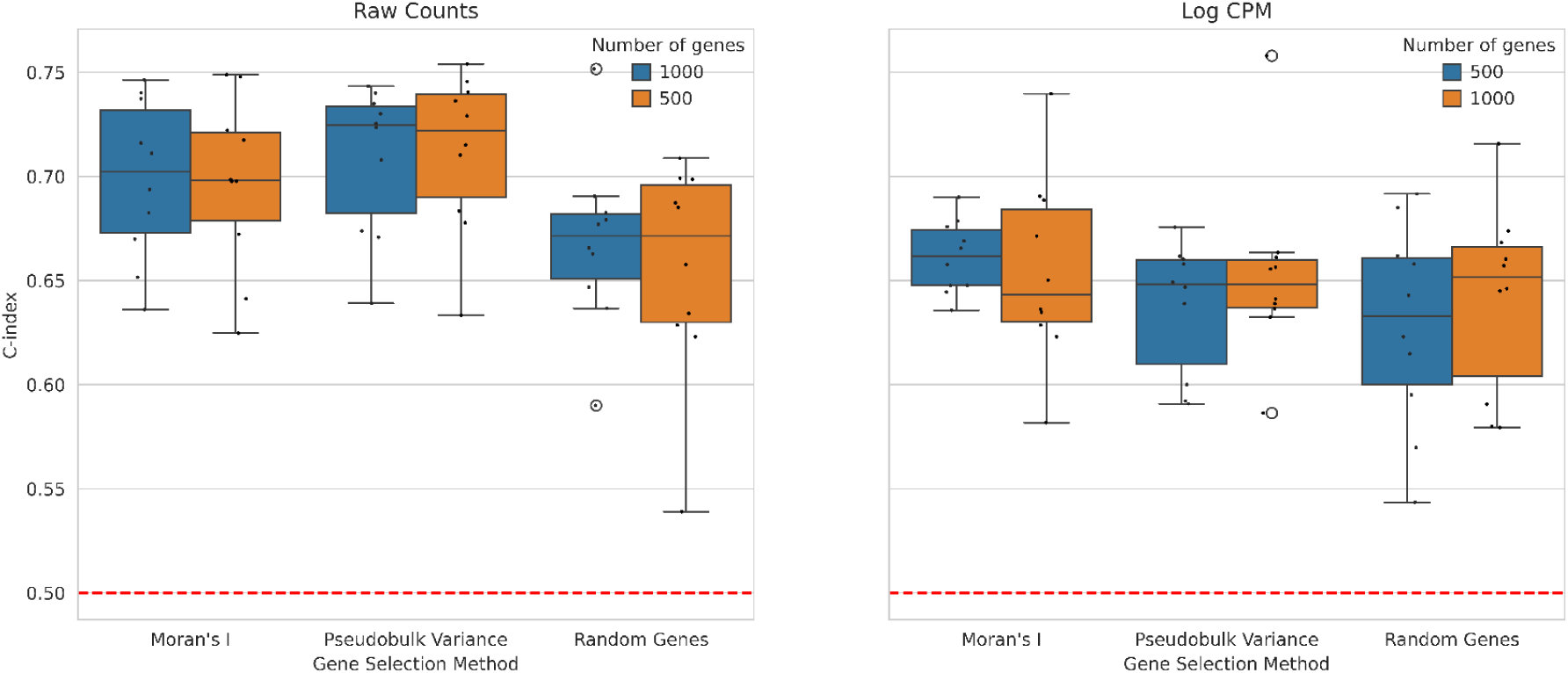
Impact of pre-processing parameter choice on model performance. The data can be left as raw counts (left) or normalised with log(1+CPM) (right). Then dimension reduction is performed by sorting genes by either pseudo-bulk variance, average Moran’s I autocorrelation, or random choice as a control. The number of genes kept after filtering is either 500 or 1000. Each boxplot represents the distribution of C-index values across repeated 5-fold cross-validation. Each dot corresponds to the mean C-index of one repeat. The red dotted line represents random chance performance.

### Supplementary Tables

**Supplementary Table S1.** Main configurations for each modality. For each modality, different configurations of the pipeline were tried. In each cell, bold denotes the best configuration, where the performance is shown in Figure 2. The H&E modality was not considered for Filtering and Scaling-normalizing. The precise number of variables filtered, the representation and prediction models hyperparameters were also tuned.

|  | Filtering | Scaling - normalizing | Representation methods | Predictive models |
| --- | --- | --- | --- | --- |
| H&E | Random sampling of tiles (training)<br>Matter detector to keep only tiles in the tissue (inference) |  | **H0 features on the whole slide**<br>H0 features aligned with SpT spots | **abMIL**<br>Chowder |
| scRNAseq | Random genes<br>**Pseudobulk most variable genes** | Raw counts<br>Log CPM counts<br>Log SCTransform<br>Raw counts (pseudobulks)<br>TPM (pseudobulks)<br>**DESeq2 (pseudobulks)** | Instances being cells:<br>Identity<br>PCA<br>scVI<br>Instances being cell type pseudobulks:<br>Identity<br>PCA<br>Pseudobulks:<br>Mean + PCA<br>**Sum + PCA** | abMIL<br>Chowder<br>**Cox** |
| bulk RNAseq | **Most variable genes** | Raw counts<br>**TPM**<br>DESeq2 | Identity<br>**PCA** | **Cox** |
| Clinical | **Set of most clinically relevant data** | **Missingness**<br>**Scaling** |  | **Cox** |
| SpT | • Random genes<br>• **Pseudobulk most variable genes**<br>• Moran's most spatially variable genes | • **Raw counts**<br>• CPM<br>• log CPM counts<br>• Raw counts (pseudobulks)<br>• TPM (pseudobulks)<br>• DESeq2 (pseudobulks) | Instances being spots:<br>• Identity<br>• **PCA**<br>• scVI<br>• scGPT-spatial<br>• Deconvolution<br>Pseudobulks:<br>• Mean + PCA<br>• Sum + PCA | • **abMIL**<br>• Chowder<br>• Cox |

**Supplementary Table S2.** Corrected resampled t-tests for all unimodal modalities. The test compares the performance of each pair of modality, ranked from worst to best predictor.

| Modality | H&E | scRNA-seq | Bulk RNA-seq | Clinical | SpT |
| --- | --- | --- | --- | --- | --- |
| H&E | / | 0.37 | 0.33 | 0.018 | 0.0061 |
| scRNA-seq | / | / | 0.43 | 0.038 | 0.014 |
| Bulk RNA-seq | / | / | / | 0.061 | 0.024 |
| Clinical | / | / | / | / | 0.32 |
| SpT | / | / | / | / | / |

**Supplementary Table S3.**
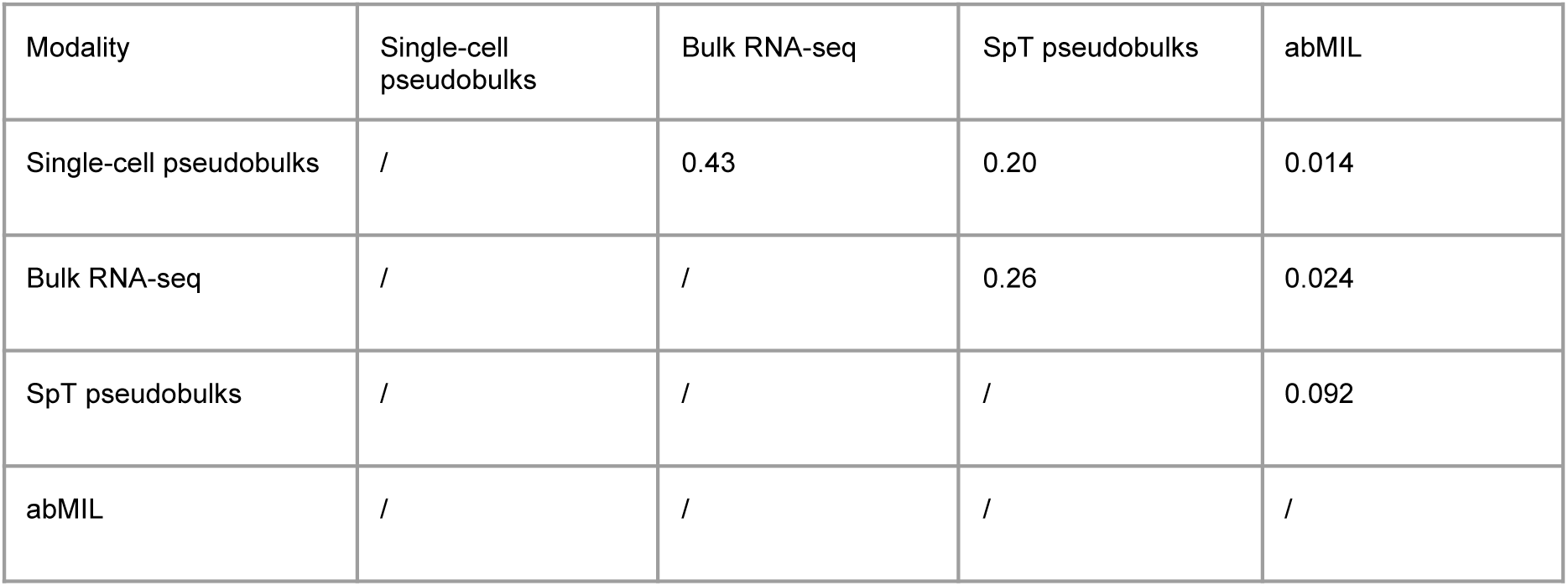
Corrected resampled t-tests for all bulk & pseudobulks. The test compares the performance of each pair of modality, ranked from worst to best predictor.

**Supplementary Table S4.** Corrected resampled t-tests for all SpT transforms. The test compares the performance of each pair of transforms, ranked from worst to best predictor.

| Spot feature extractor | SpT counts (no transform) | scVI | scGPT-spatial | Deconvolution | PCA |
| --- | --- | --- | --- | --- | --- |
| SpT counts (no transform) | / | 0.37 | 0.16 | 0.054 | 0.028 |
| scVI | / | / | 0.28 | 0.12 | 0.064 |
| scGPT-spatial | / | / | / | 0.28 | 0.17 |
| Deconvolution | / | / | / | / | 0.36 |
| PCA | / | / | / | / | / |

**Supplementary Table S5.** Batch effect assessment. The “label” is the batch effect assessment, here the sample effect. The biological conservation is measured with the dominant cell type in every spot. Both avg_bio and avg_batch are between 0 and 1, the closer to 1 meaning a better biological conservation of the majority cell type and a better sample mixing, respectively.

|  | No transform | scVI | scGPT - spatial | Deconvolution | PCA |
| --- | --- | --- | --- | --- | --- |
| NMI_cluster/<br>label | 0.14 | 0.13 | 0.19 | 0.25 | 0.14 |
| ARI_cluster/<br>label | 0.0091 | 0.0085 | 0.037 | 0.10 | 0.0089 |
| ASW_label | 0.32 | 0.51 | 0.53 | 0.45 | 0.35 |
| ASW_label/<br>batch | 0.71 | 0.90 | 0.80 | 0.69 | 0.72 |
| graph_conn | 0.61 | 0.68 | 0.90 | 0.94 | 0.45 |
| **avg_bio** | **0.16** | **0.21** | **0.25** | **0.26** | **0.17** |
| **avg_batch** | **0.66** | **0.79** | **0.85** | **0.82** | **0.59** |

**Supplementary Table S6.** Corrected resampled t-tests for H&E and SpT+H&E modalities. The test compares the performance of each pair of modality, ranked from worst to best predictor.

| Modality | H&E-only | MabMIL - HE-only<br>inference | MabMIL - multimodal<br>inference |
| --- | --- | --- | --- |
| H&E-only | / | 0.32 | 0.011 |
| MabMIL - HE-only inference | / | / | 0.036 |
| MabMIL - multimodal inference | / | / | / |

## Extended data

**Extended Data Table 1.**
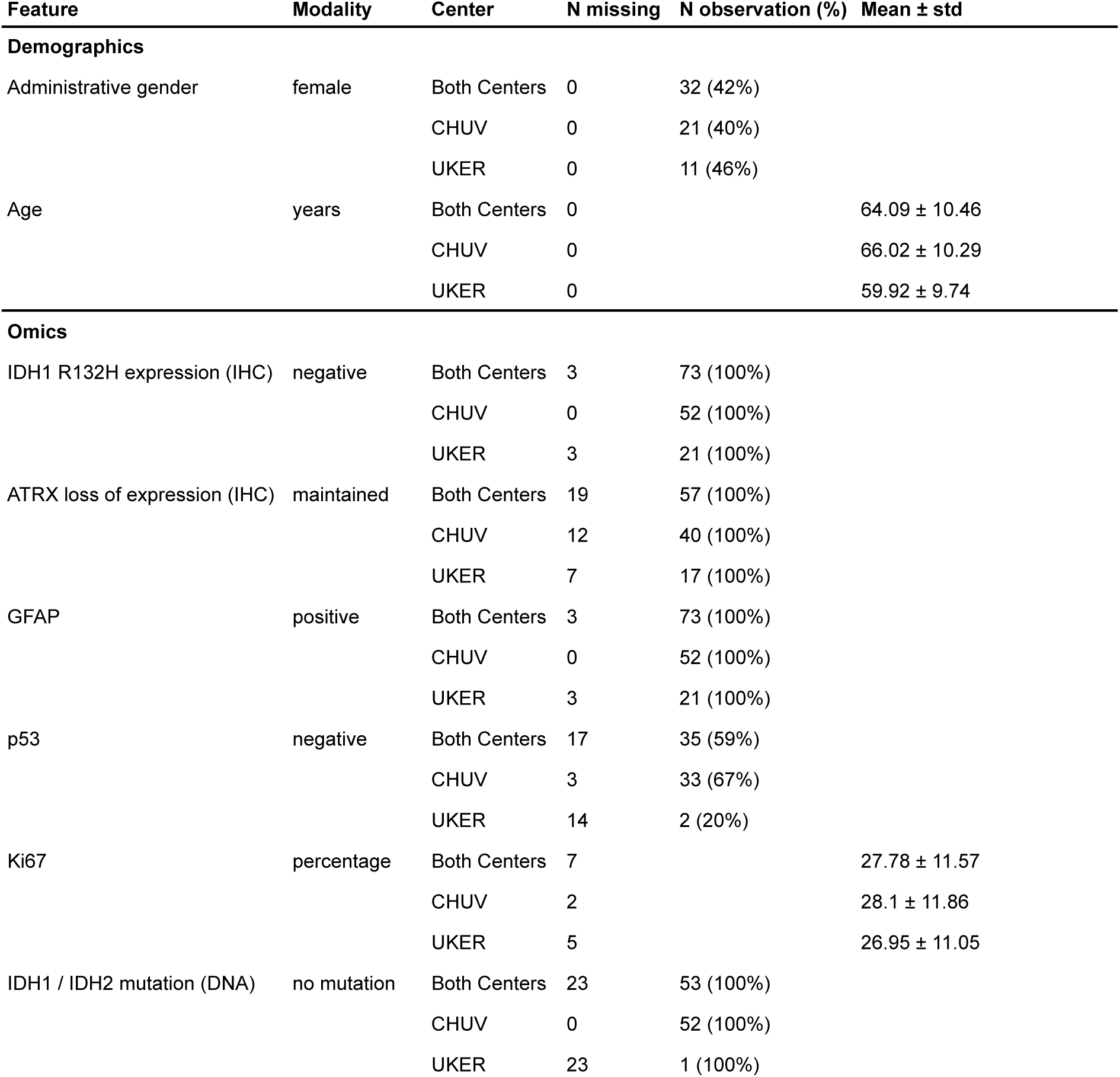
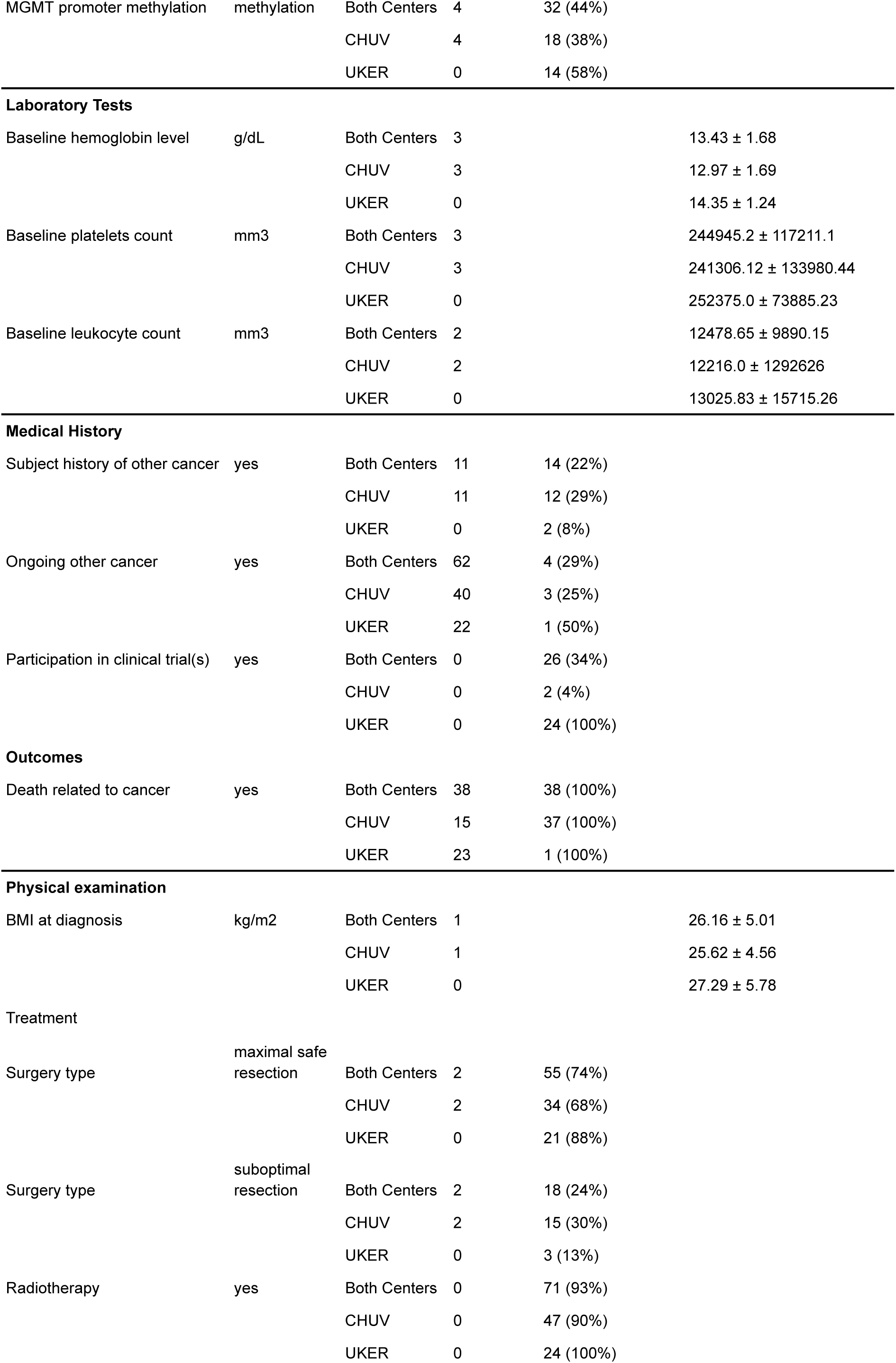
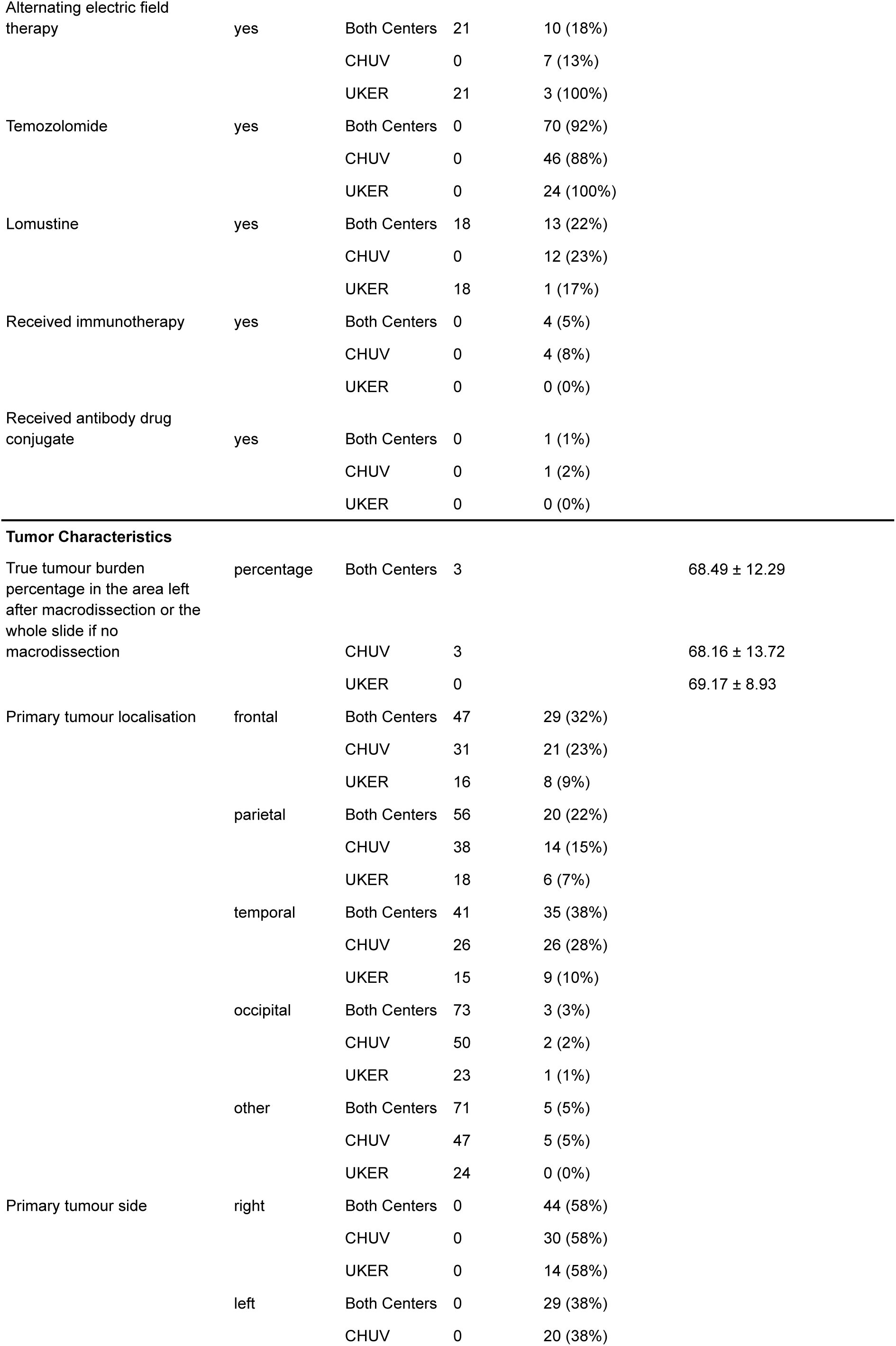
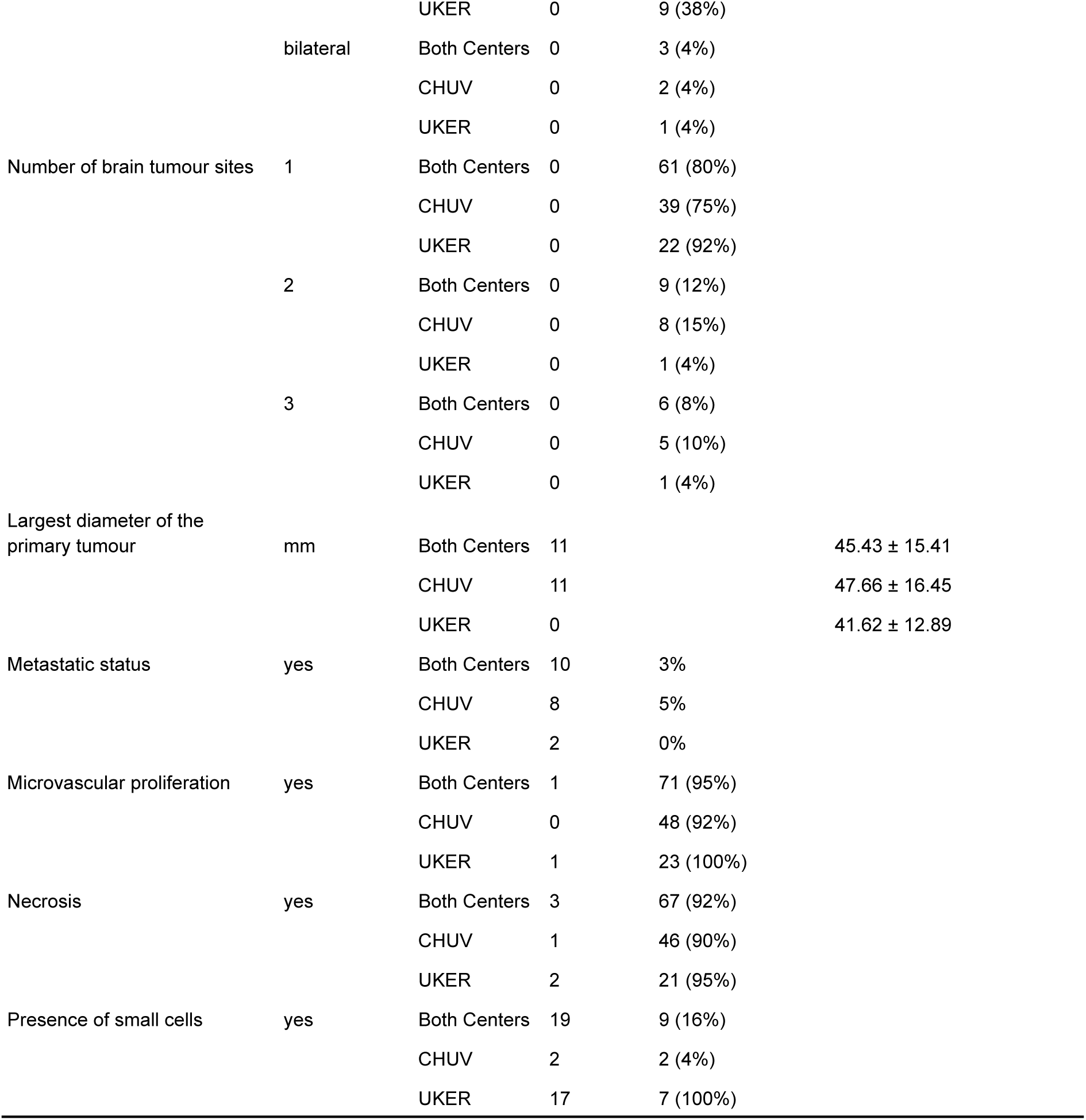
Cohort Characterization.

## Code availability

The code to replicate our results will be made publicly available upon acceptance of the paper.

## Acknowledgements

We are grateful to patients, physicians, nurses and research assistants involved in the study. This study makes use of a glioblastoma cohort of the MOSAIC consortium (Owkin; Charité – Universitätsmedizin Berlin (DE); Lausanne University Hospital - CHUV (CH); Universitätsklinikum Erlangen (DE); Institut Gustave Roussy (FR); University of Pittsburgh (USA)), a non-interventional clinical trial registered under NCT06625203.

